# The fastest movements and the most extreme deformations of the cell nucleus are driven by microtubules in the model fungus *Podospora anserina*

**DOI:** 10.1101/2025.06.08.658251

**Authors:** Viron Marie-caroline, Kachaner Zoé, Grognet Pierre, Lalanne Christophe, Guichet Antoine, Brun Sylvain

## Abstract

A growing number of studies shows that nuclear movement, positioning, and the mechanical constraints applied to the nucleus play important roles in cellular physiology and gene regulation. This process, known as nuclear mechanotransduction, is an emerging field of research, and its deregulation can lead to severe human pathologies. In this study, we examine the behavior of female and male nuclei during sexual reproduction in the model fungus *Podospora anserina*. After plasmogamy, the two types of nuclei do not immediately undergo karyogamy. Instead, once inside the trichogyne (the female-specific hyphae extending from the ascogonium) the male nuclei migrate over distances of up to several hundred micrometers from the trichogyne tip, where plasmogamy occurs, to the ascogonium at the center of the protoperithecium. Our study highlights the contrasting behaviors of the two nuclear types: female nuclei remain immobile and spherical, while male nuclei move rapidly and exhibit striking deformations and stretching. Trichogynes are branched and compartmentalized, consisting of articles separated by septa that communicate through the septal pores. We show that during their migration across these compartments, male nuclei undergo the most extreme movements, deformations, and speeds reported to date for nuclei in motion. We also investigated the role of the actin and microtubule (MT) cytoskeletons in male nuclei behavior. Both cytoskeletal networks are highly dynamic and abundant in trichogynes, particularly at septa. We demonstrate that while actin depolymerization does not impair male nuclei migration, MT disruption causes a complete arrest of both nuclear movement and stretching, indicating a critical role for MTs. Finally, we reveal that the MT network in trichogynes is highly polarized, offering new insights into how male nuclei navigate within branched trichogynes.

## Introduction

The study of the cell nucleus has always been central to biology. Research into cell signaling and gene expression regulation has been pivotal in understanding how cells function, how they perceive their environment, how tissues organize, how organisms develop, and how diseases arise from impaired signaling or disrupted gene expression. In comparison, the understanding of mechanotransduction—that is, how physical forces and constraints applied to the nucleus can alter its organization and change gene expression—has only recently emerged as a new and growing field of research (Dahl et al., 2008; Graham and Burridge, 2016; Kirby and Lammerding, 2018; Maurer and Lammerding, 2019; Janota et al., 2020). The idea that signals transmitted to the nucleus could be mechanical in nature was largely ignored for a long time. However, an increasing number of human diseases are now being recognized as resulting primarily from failures in mechanotransduction (Dauer and Worman, 2009; Davidson and Lammerding, 2014). Similarly, defective nuclear positioning has been linked to severe disorders such as microcephaly and myopathies (Dauer and Worman, 2009; Alvarado-Kristensson and Rosselló, 2019). Furthermore, abnormalities in nuclear envelope (NE) stiffness are associated with serious pathologies such as progeria and cancer (Davidson and Lammerding, 2014; Denais and Lammerding, 2014). Interestingly, it has been reported that reducing microtubule (MT)-generated forces on the NE can rescue defects associated with laminopathies (Larrieu et al., 2014). Nuclear deformation also plays a critical role in cancer cell dissemination. Because the nucleus is the largest and stiffest organelle in the cell, it becomes a limiting factor during cancer cell migration through tissue (Infante et al., 2018). The nuclei of metastatic cells undergo extreme deformation as they squeeze through tight spaces when penetrating or exiting the blood vessels for instance. Recent studies have shown that repeated nuclear deformation affects NE stiffness, chromatin organization, and gene expression, contributing to increased tumor aggressiveness (Golloshi et al., 2022). Mechanotransduction and nuclear movement have been studied in a variety of model organisms, ranging from yeast and Drosophila to humans. These studies have shown that nuclei interact with various cytoskeletal elements, and that their movements are dependent on them. Unsurprisingly, MTs, actin, and their associated motor proteins are involved in both nuclear movement and mechanotransduction (Janota et al., 2020; Xiang, 2018; Lepesant et al., 2024; Tissot et al., 2017). Lamins which play a crucial role in nuclei stiffness are also involved in these processes (Davidson and Lammerding, 2014); however, lamins have not been identified in fungi, raising questions about how fungal nuclei resist mechanical constraints.

In filamentous fungi, nuclear movement was first studied in vegetative hyphae (Gladfelter and Berman, 2009; Xiang, 2018) and more recently during sexual reproduction in the model fungus *Neurospora crassa* (Brun et al., 2021). Here, we investigate this process in the model fungus *Podospora anserina*, a well-established organism for genetic and cell biology analysis (Silar, 2013). Its short life cycle (7 days), the production of small, easily imaged fruiting bodies, and the ease of generating transgenic lines facilitated our cell biology and genetics approaches. In *P. anserina,* sexual reproduction between individuals of opposite mating types (heterothallic cross) is initiated when “female” trichogynes are attracted to pheromones secreted by “male” spores (spermatia in *P. anserina* and conidia in *N. crassa*) as in *N. crassa* (Kim et al., 2012). Eventually, the two structures fuse which represents the fertilization/plasmogamy step *per se*. Trichogynes are specialized sexual hyphae emitted by ascogonia, the female structures embedded in developing protoperithecia (fruiting bodies) (Fig 1). These trichogynes, which can extend over hundreds of micrometers, are composed of successive compartments (called articles) separated by septa and connected via pores. Trichogynes contain female nuclei and, importantly after plasmogamy, the male nuclei that enter the trichogynes do not fuse with female nuclei but migrate toward the ascogonia (Bennett and Turgeon, 2016).

**Fig 1:**
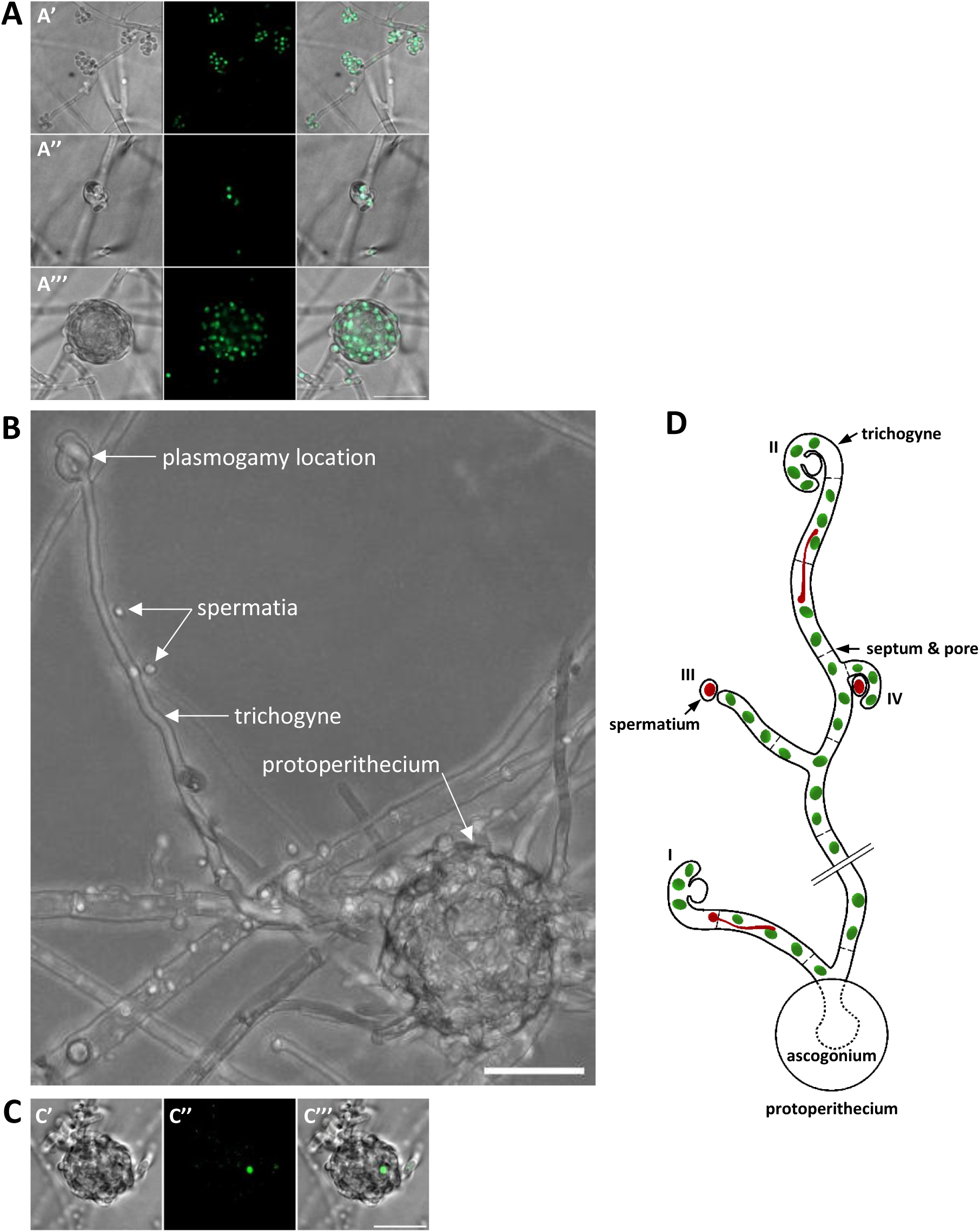
Reproductive organs engaged in sexual reproduction in *P. anserina*. **A)** *H1-GFP* strain. First row: conidiophores bearing spermatia; second row: ascogonium; third row: protoperithecium. Images are shown from left to right: transmitted light, GFP fluorescence, and merged channels. **B)** Z-projection image of a protoperithecium undergoing fertilization. The spermatium, bound by the trichogyne, is concealed by the trichogyne itself and is not visible in the image. **C)** Wild-type protoperithecium fertilized by an H1-GFP-labeled male nucleus. Images from left to right: transmitted light, GFP fluorescence, and merged channels. Scale bars: 20 µm. **D)** Schematic representation of the fertilization process in *P. anserina* using a cross between male *H1-mCherry* and female *H1-GFP* strains. The ascogonium, the “female” cell, lies at the core of the protoperithecium and produces trichogynes—sex-specific hyphae guided by pheromones released by “male” spermatia of the opposite mating type (Peraza-Reyes and Malagnac, 2016; Kim et al., 2012). Spermatia contain a single nucleus. Trichogynes consist of a series of compartments (articles) separated by septa with septal pores that allow cytoplasmic continuity. Female nuclei (green) are distributed along the trichogyne and may undergo mitosis during trichogyne growth. We distinguish four modes of spermatium binding by trichogynes: **Type I:** binding by a newly growing trichogyne emerging from the protoperithecium, **Type II:** binding by the main apex of a trichogyne already extended on the medium at the time of spermatium inoculation, **Type III:** binding by a trichogyne branch specifically attracted to the spermatium, **Type IV:** binding by a small hook-like protrusion emerging from the trichogyne. After plasmogamy, male and female nuclei do not immediately fuse (karyogamy). Instead, male nuclei (red), contributed by spermatia attached at various points, migrate up the trichogyne toward the ascogonium. During this migration, male nuclei undergo significant deformation and elongation as they navigate through septal pores and bypass female nuclei.

Here, we provide a detailed description of the fertilization process, as well as of male nuclei movements and deformations during migration in *P. anserina*. During their migration, male nuclei exhibit an “inchworm-like” movement—repeatedly contracting and stretching as they bypass female nuclei or pass-through septal pores (Fig 1D). Our study shows that the overall process is conserved compared to *N. crassa*, but we observed notable differences, including the potential for polyfertilization of protoperithecia and more dramatic speeds and deformations of male nuclei in *P. anserina*. We also investigated the actin and MT networks in trichogynes and their respective roles in male nuclei migration. Finally, we demonstrate that the MT network in trichogynes is highly polarized and that the remarkable movements and deformations of male nuclei are dependent on MTs.

## Results

### The male and female structures involved in sexual reproduction

To gain insights into the complete fertilization process in *Podospora anserina*, we performed live imaging of fertilization at low magnification (25x or 40x objectives), collecting data throughout all stages—from trichogyne growth to the eventual arrival of male nuclei in the protoperithecia. The fungal strains used in our experiments were genetically homogeneous, with all nuclei within an individual fungal thallus being genetically identical (homokaryotic). These homokaryotic *P. anserina* strains are hermaphroditic (Fig 1), differentiating both “female” ascogonia (Fig 1A’’) and “male” spermatia (Fig 1A’). Spermatia are mitospores similar to conidia; they are produced by conidiophores but, unlike typical conidia, cannot germinate. Instead, they function exclusively as “male” donor cells during fertilization (Peraza-Reyes and Malagnac, 2016). Ascogonia initially appear as twisted hyphal branches a few micrometers in length and are rapidly surrounded by adjacent vegetative hyphae (maternal tissue), forming protoperithecia (Fig 1A’’’, 1C) (Zickler et al., 1995). While the protoperithecia gradually expand to reach diameters of several tens of micrometers, the ascogonia embedded within them produce trichogynes. In the absence of fertilization by spermatia of the opposite mating type, these trichogynes continue to grow and branch (Figs 1B, 1D). Consequently, the 5- to 7-day-old thalli used in our study exhibited a fertile female network interwoven with vegetative hyphae. This fertile network, composed of trichogynes emerging from ascogonia, can only be fertilized by spermatia of the opposite mating type. *P. anserina* homokaryotic strains possess a single mating type—either *mat−* (*MAT-1-1*) or *mat+* (*MAT-1-2*)—and sexual reproduction occurs exclusively between strains of opposite mating types (heterothallism). In our experiments, we used *mat+* spermatia to fertilize *mat−* female mycelium, and conversely, *mat−* spermatia to fertilize *mat+* mycelium. We did not observe any difference between the two reciprocal crosses. Therefore, for the sake of simplicity, the mating types of the strains used are not specified further in the manuscript.

### Trichogyne chemotropic growth and spermatium binding

In our experiments, we tracked male nuclei inside spermatia or during their migration, as well as female nuclei inside trichogynes, by tagging nuclear chromatin with GFP-tagged (H1-GFP) and mCherry-tagged (H1-mCherry) histone H1 (see Materials and Methods, Table 1). When spermatia were inoculated onto the “female” thallus (also referred to as the female partner or mycelium) of the opposite mating type, we observed trichogynes (or their branches) growing toward the spermatia in an undulating manner reminiscent of chemoattraction, at a speed of 0.6 ± 0.3 µm/min (Figs 1B, 1D, 2; Movie 1; Table S1). Movie 1 illustrates the growth of a trichogyne eventually binding a spermatium (circled in Fig 2), culminating in the migration of the male nucleus up the trichogyne (Fig 2). As commonly observed, the trichogyne pushed and displaced the spermatium by a few micrometers after binding (t = 76 min). Seventy-two minutes later (t = 148 min), we observed the entry of the male nucleus expressing H1-mCherry into the trichogyne, followed by its migration toward the protoperithecium. As shown in Movie 1, the female nuclei residing within the trichogyne could undergo mitosis during trichogyne growth. In contrast, we never observed mitosis of female nuclei in trichogynes that had ceased growing due to spermatium binding.

**Fig 2.**
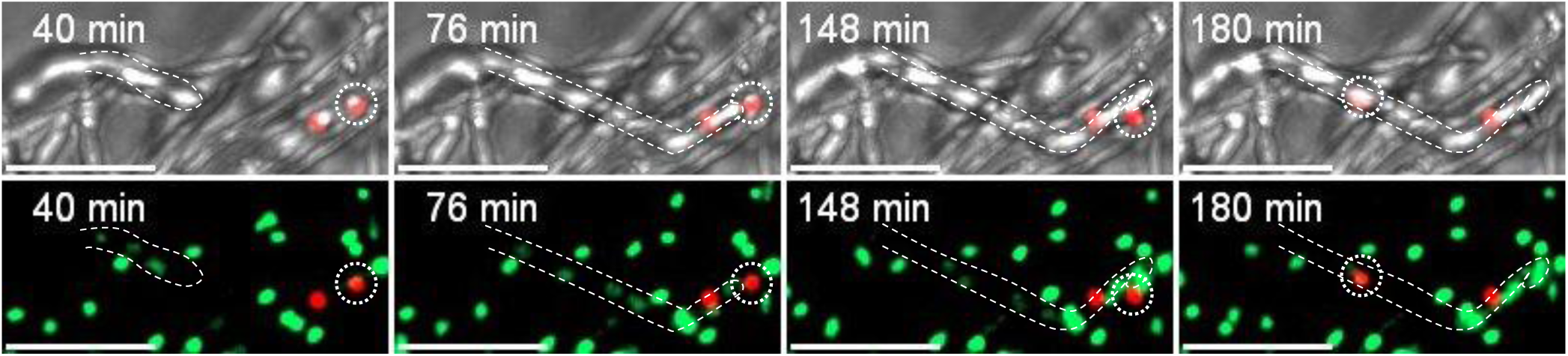
Trichogyne chemotropic growth followed by plasmogamy and male nucleus migration. Cross: male *H1-mCHerry* X female *H1-GFP*. Z-projection images from time-lapse movie 1 acquired using a spinning disk microscope (1 frame every 4 minutes). In the first row, transmitted light and mCherry channels are merged to visualize H1-mCherry-labeled male spermatia and nuclei (red). In the second row, GFP and mCherry channels are merged to simultaneously visualize H1-GFP-labeled female nuclei (green) and H1-mCherry-labeled male nuclei (red). The growing trichogyne is outlined with a dashed line, and the spermatium attracting the trichogyne is circled with a dashed line. At t = 76 min, the trichogyne establishes contact with the H1-mCherry-labeled spermatium. At t = 148 min, the H1-mCherry-labeled male nucleus has just entered the trichogyne. At t = 180 min, the H1-mCherry-labeled male nucleus (dashed circle) is migrating along the trichogyne toward the protoperithecium, located just outside the left edge of the field of view. Scale bar: 20 µm.

**Table 1:**
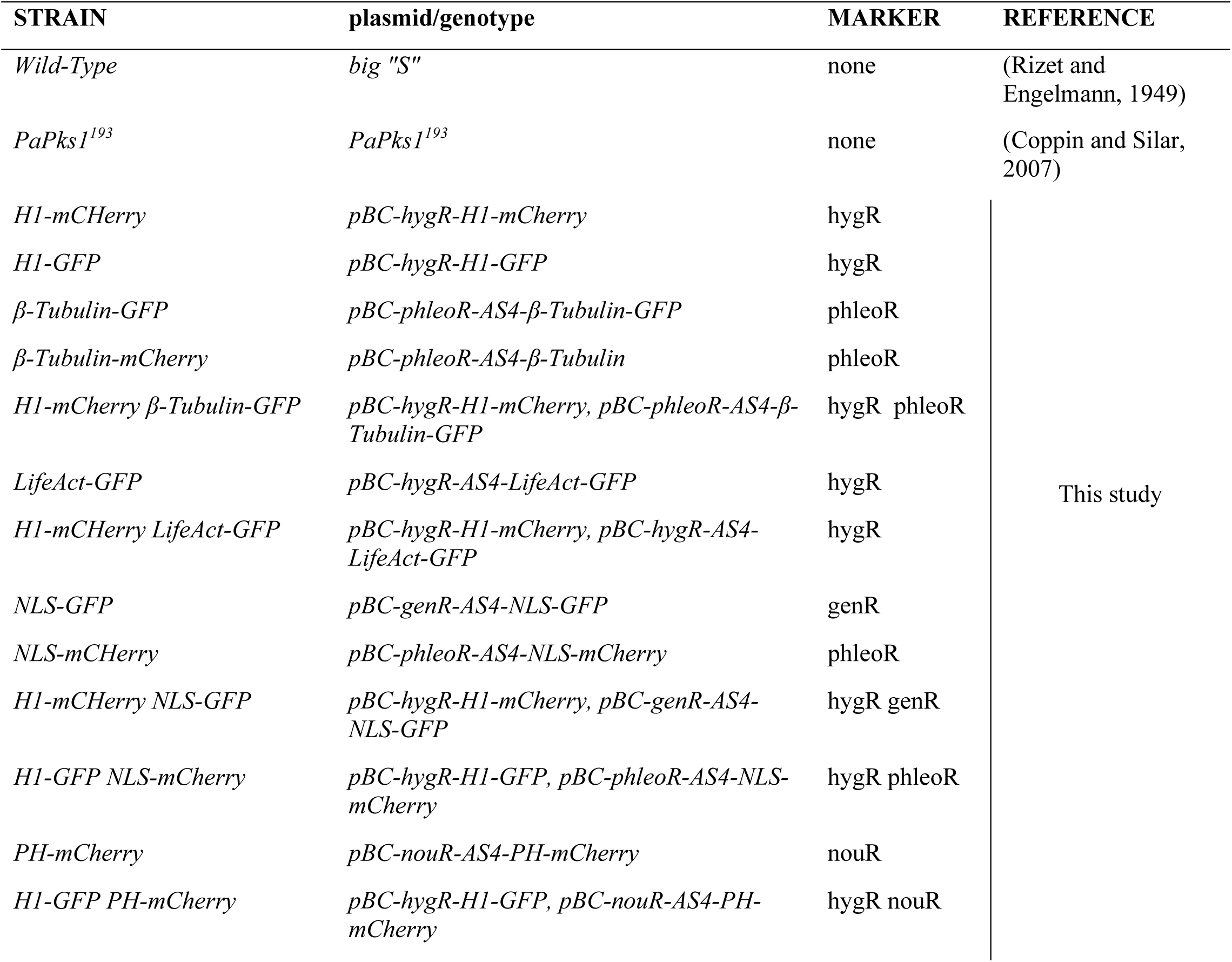

| STRAIN | plasmid/genotype | MARKER | REFERENCE |
| --- | --- | --- | --- |
| *Wild-Type* | *big "S"* | none | (Rizet and Engelmann, 1949) |
| *PaPks1*<sup>193</sup> | *PaPks1*<sup>193</sup> | none | (Coppin and Silar, 2007) |
| *H1-mCherry* | *pBC-hygR-H1-mCherry* | hygR | This study |
| *H1-GFP* | *pBC-hygR-H1-GFP* | hygR |  |
| *β-Tubulin-GFP* | *pBC-phleoR-AS4-β-Tubulin-GFP* | phleoR |  |
| *β-Tubulin-mCherry* | *pBC-phleoR-AS4-β-Tubulin* | phleoR |  |
| *H1-mCherry β-Tubulin-GFP* | *pBC-hygR-H1-mCherry, pBC-phleoR-AS4-β-Tubulin-GFP* | hygR phleoR |  |
| *LifeAct-GFP* | *pBC-hygR-AS4-LifeAct-GFP* | hygR |  |
| *H1-mCherry LifeAct-GFP* | *pBC-hygR-H1-mCherry, pBC-hygR-AS4-LifeAct-GFP* | hygR |  |
| *NLS-GFP* | *pBC-genR-AS4-NLS-GFP* | genR |  |
| *NLS-mCherry* | *pBC-phleoR-AS4-NLS-mCherry* | phleoR |  |
| *H1-mCherry NLS-GFP* | *pBC-hygR-H1-mCherry, pBC-genR-AS4-NLS-GFP* | hygR genR |  |
| *H1-GFP NLS-mCherry* | *pBC-hygR-H1-GFP, pBC-phleoR-AS4-NLS-mCherry* | hygR phleoR |  |
| *PH-mCherry* | *pBC-nouR-AS4-PH-mCherry* | nouR |  |
| *H1-GFP PH-mCherry* | *pBC-hygR-H1-GFP, pBC-nouR-AS4-PH-mCherry* | hygR nouR |  |

We observed that trichogynes could bind to spermatia in different ways. Spermatia could be bound by i) the apex of newly emerging trichogynes projecting directly from protoperithecia (Type I), ii) the principal apex of pre-existing trichogynes on the thallus at the time of inoculation (Type II), iii) the apex of branches that developed on trichogynes and growing toward spermatia (Type III), or iv) small hyphal hooks emerging from trichogynes and binding spermatia already in contact with the trichogyne (Type IV) (Fig 1D). Importantly, we never observed spermatium/trichogyne binding involving trichogynes growing from young, bare ascogonia. Fertilization was only observed between spermatia and trichogynes emanating from properly formed protoperithecia. We further determined that the smallest protoperithecia fertilized by male nuclei had a diameter of 21 µm.

### Plasmogamy

The initial step of plasmogamy described above—namely, the binding of trichogynes to spermatia—occurred serendipitously on female thalli following the inoculation of spermatia of the opposite mating type. To image the entire fertilization process, live—from trichogyne growth to the arrival of male nuclei in protoperithecia—while addressing its inherent unpredictability, we conducted live-recording sessions at low magnification. These recordings captured both protoperithecia and spermatia within the field of view over extended periods (10 to 15 hours), using spermatia expressing H1-mCherry or H1-GFP. This protocol enabled us to document the complete fertilization process as previously described and to collect valuable data on its various stages. Notably, we determined that the average time between trichogyne–spermatium binding and the entry of the male nucleus into the trichogyne was 89 ± 33 minutes (Table S1). However, under these experimental conditions, we were unable to directly observe the cell fusion event that precedes male nuclei entry into the trichogyne. To address this limitation, we used spermatia doubly tagged with H1-mCherry and NLS-GFP or H1-GFP and NLS-mCherry. In these spermatia, chromatin is labeled with H1-mCherry or H1-GFP, while both the nucleoplasm and cytoplasm are tagged with NLS-GFP or NLS-mCherry (*i.e*., GFP or mCherry fused to a type I Nuclear Localization Signal, NLS). Although NLS-GFP and NLS-mCherry primarily accumulated in the nuclei, the cytoplasmic fluorescence of both markers was sufficiently strong for detection (data not shown). Following inoculation of these double-tagged spermatia, we tracked male nuclei before and during migration based solely on H1-mCherry or H1-GFP chromatin signals. Importantly, we observed that the fluorescent signal corresponding to NLS-GFP or NLS-mCherry disappeared from the spermatia ∼3 min, as an average, before the onset of male nuclei migration (Fig 3, Movie 2, Table S1). We interpreted this loss of fluorescence as a result of the immediate dilution of spermatial cytoplasmic content into the trichogyne cytoplasm upon cell fusion. Thus, the disappearance of the NLS-GFP/NLS-mCherry signal likely marked the precise moment of cell fusion between spermatia and trichogynes. Accordingly, male nuclei entry into the trichogynes occurs rapidly after fusion, especially when compared to the ∼90-minutes interval following initial trichogyne–spermatium binding.

**Fig 3.**
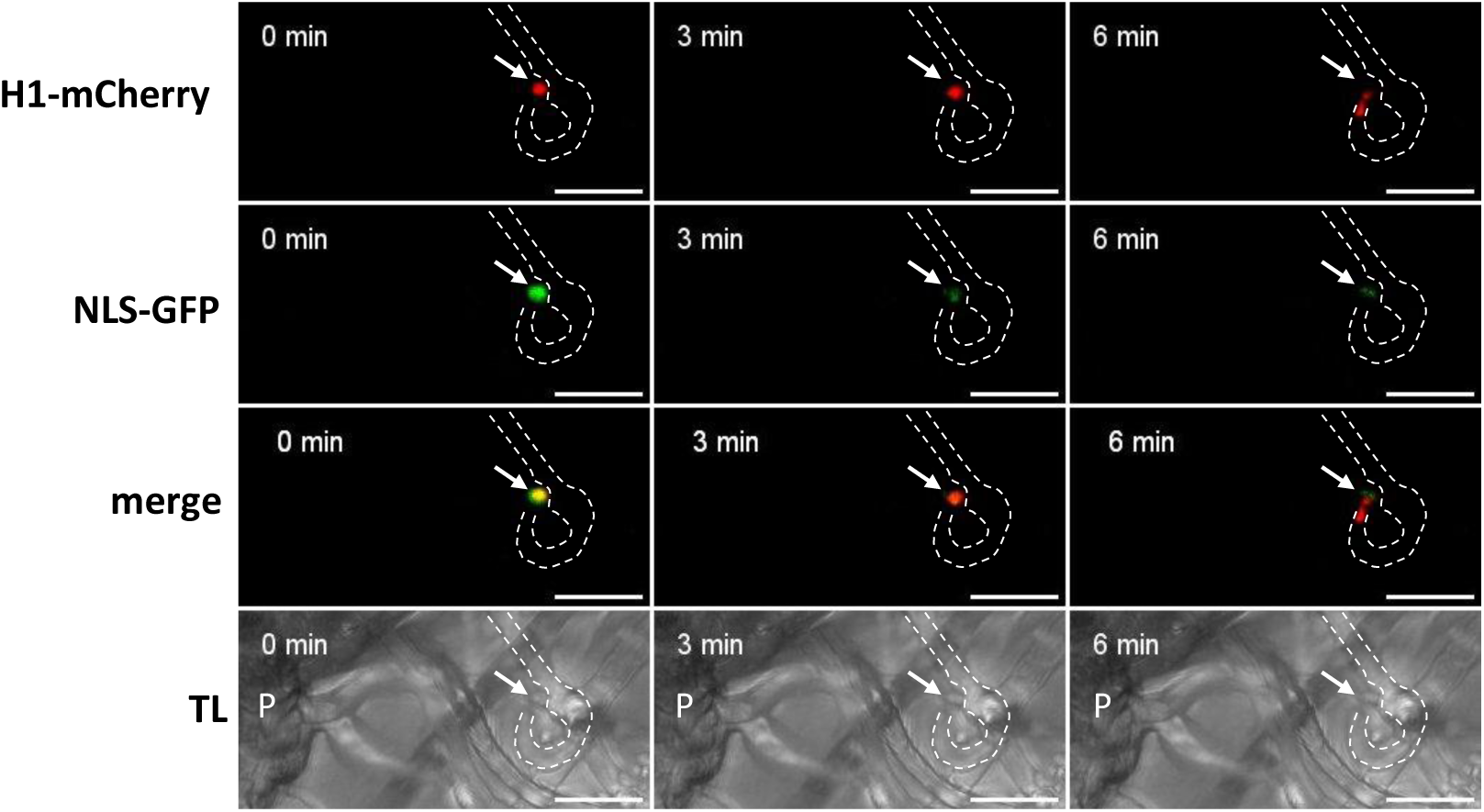
Plasmogamy followed by male nucleus entry into the trichogyne. Cross: male *H1-mCherry NLS-GFP* X female *wild-type*. Z-projection images from time-lapse movie 2 acquired using a spinning disk microscope (1 frame every 3 minutes). The trichogyne is outlined with a dashed line, and the spermatium is indicated by an arrow. At t = 0 min, both NLS-GFP and H1-mCherry signals are visible in the spermatium. At t = 3 min, the NLS-GFP signal disappears, indicating occurrence of plasmogamy. At t = 6 min, the H1-mCherry-labeled male nucleus begins to enter the trichogyne. TL: transmitted light; P: protoperithecium. Scale bar: 10 µm.

### Polyfertilization of perithecia

As previously mentioned, trichogynes are often branched, allowing them to fuse with multiple spermatia. This likely leaded to the entry and migration of several male nuclei, which may ultimately reach and enter the same protoperithecium (Fig 1D). To quantify the fertilization rate of protoperithecia and assess the occurrence of polyfertilization, we conducted live imaging of the fertilization process (male *H1-mCherry* X female *WT*) (wild-type) over a 15-hour period. Recordings were performed at low magnification (25× objective), beginning 4 hours after spermatia inoculation. We observed that 70% of the recorded protoperithecia (16 out of 23) were fertilized, and notably, half of these (8 out of 16) were fertilized by more than one nucleus during the time-lapse (Fig S1A, Movie S1). In Movie S1, three male nuclei entered the same protoperithecium (male *H1-mCHerry* X female *H1-GFP* cross). In one case, we observed five nuclei entering a single protoperithecium (data not shown). This observation raised the question of whether fertilization by spermatia carrying different genotypes could result in perithecia with mosaic contents, that is producing asci of different paternal origin. To test this hypothesis, we fertilized the wild-type *S* strain (which produces dark melanized ascospores) with a mixture of wild-type spermatia and spermatia from the *PaPks1^193^* mutant (which produces unpigmented, non-melanized ascospores) (Coppin and Silar, 2007). We dissected 158 perithecia and found the following: 102 perithecia contained only asci with four melanized ascospores, indicating fertilization by wild-type spermatia. 56 perithecia contained asci with two melanized and two non-melanized ascospores, consistent with fertilization by *PaPks1^193^*spermatia. No perithecia displayed a mosaic composition containing both types of asci (Fig S1B). These results indicates that, despite frequent polyfertilization events at the protoperithecium level, mature perithecia give rise to asci derived from a single paternal genotype.

### Difference between female and male nuclei behavior in trichogynes

We crossed male and female strains expressing differentially tagged nuclei to enable the independent tracking of each nuclear type during fertilization. Notably, in both male *H1-mCherry* X female *H1-GFP* and male *H1-GFP* X female *H1-mCherry* crosses, we never observed any merging or overlap of the green and the red fluorescence signal within the nuclei. This allowed the clear and simultaneous observation of male and female nuclei within the same trichogynes. We first characterized the behavior of female nuclei in trichogynes, both before plasmogamy and after plasmogamy, during male nuclei migration. Prior to plasmogamy, female nuclei in trichogynes: i) were distributed along the entire length of the trichogynes, ii) were round with an average diameter of 2.1 ± 0.2 µm, iii) exhibited oscillatory movements, and iv) underwent division exclusively during trichogyne growth (Table S1; Figs 1D, 2, 4, and 5; Movies 1, 3, and 4). After spermatium binding, female nuclei within the terminal article of the trichogyne tended to accumulate near the tip. Following plasmogamy, both male and female nuclei coexisted within the trichogyne: male nuclei migrated from the spermatia toward the protoperithecia, while female nuclei generally remained stationary. Interestingly, female nuclei in trichogynes undergoing male nuclei migration displayed a significant reduction in oscillatory speed compared to their behavior before plasmogamy (Fig 4). Unlike the predominantly round female nuclei, male nuclei were never round during migration. Instead, they displayed remarkable morphological plasticity, transitioning between relaxed and extremely elongated forms. In their relaxed state, male nuclei had a mean length of 6.2 ± 1.0 µm (Table S1), whereas the maximum observed length for a stretched male nucleus reached up to 71 µm (Fig S2, Movie S2).

**Fig 4.**
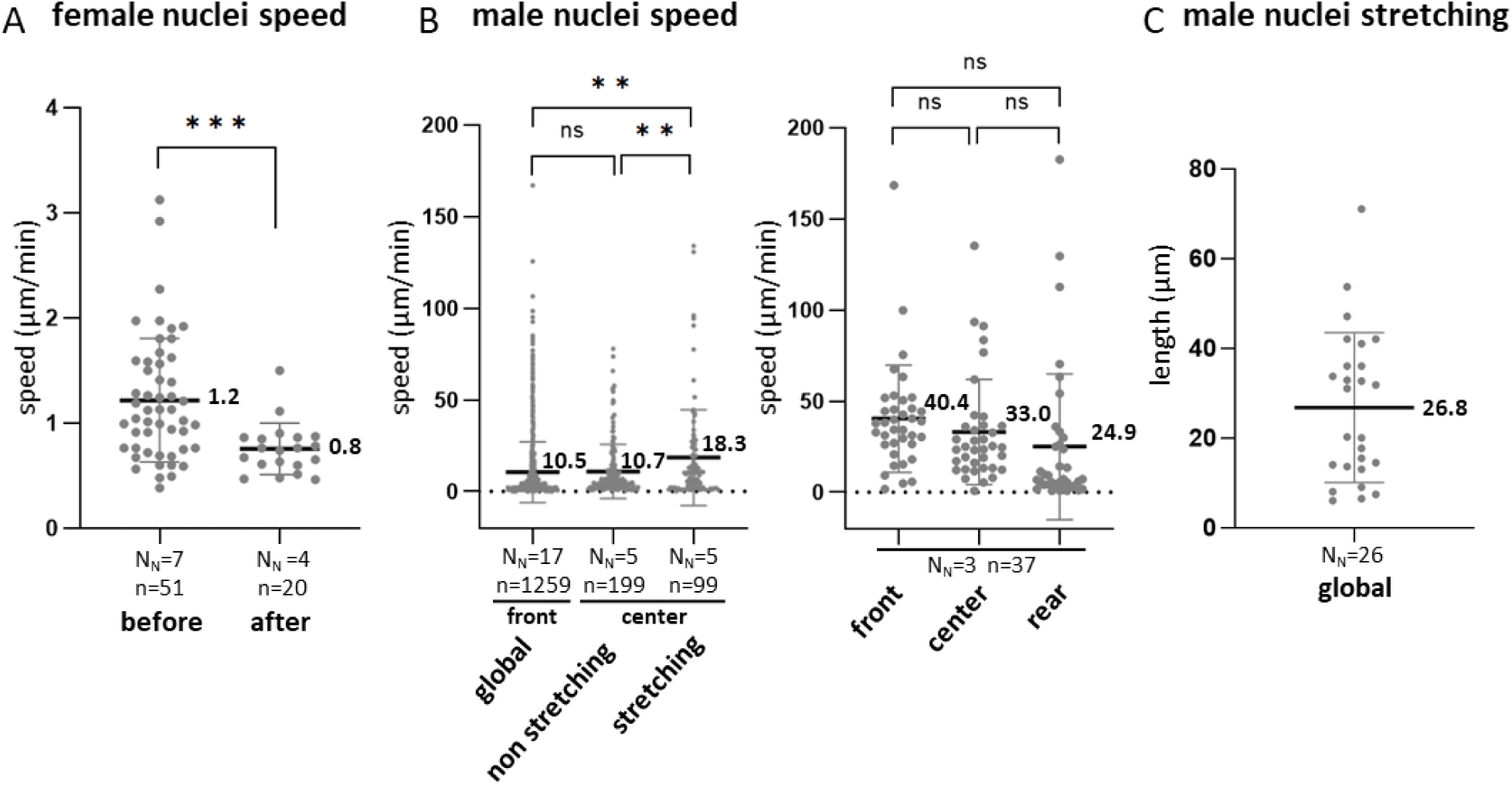
Speeds of female and male nuclei and male nuclei stretching during migration. **A)** Cross: male *H1-mCherry* X female *H1-GFP*. Comparison of female nuclei speed before plasmogamy and after plasmogamy, during the migration of male nuclei; mean speeds are indicated on the graph. **B)** Global: compilation of male nuclei front speed measurements from several crosses: male *H1-GFP* X female *H1-mCherry*, male *H1-mCherry* X female *β-Tubulin-GFP*, male *H1-GFP* X female *PH-mCherry*, and male *H1-mCherry β-Tubulin-GFP* X female *wild-type*. Non-stretching and stretching: center speed measurements of male nuclei from *H1-mCherry* X female *β-Tubulin-GFP* crosses, divided into two categories—non-stretching: time points where the nuclei were in motion without stretching; stretching: time points showing active stretching. Front, center, and rear speed measurements are compiled from crosses involving male *H1-GFP* X female *PH-mCherry*, male *H1-mCherry* X female *LifeAct-GFP*, and male *H1-mCherry* X female *β-Tubulin-GFP* (see also Fig S3). **C)** Global: compilation of the maximum stretching length of male nuclei from crosses including *H1-mCherry* X female *β-Tubulin-GFP*, male *H1-GFP* X female *PH-mCherry* and male *H1-mCherry β-Tubulin-GFP* X female *wild-type*. N_N_: number of nuclei analyzed; n: number of individual speed measurements. Statistical analysis: Welch’s t-test. P-values: ns = not significant; * P < 0.05; ** P < 0.01; *** P < 0.001; **** P < 0.0001.

### Male nuclei migration

Male nuclei migration was a highly complex and discontinuous process, characterized by phases of acceleration, periods of stalling, back-and-forth movements, changes of direction at branch points, and notably, phases of extreme nuclear stretching. To accurately describe these movements, we assigned each male nucleus a front, center, and rear, and analyzed their respective movements and speeds separately during migration (Figs 4, 5, and S3). Our primary focus was on the front speed when describing male nuclei migration, but we also compared speeds at the front, center, and rear to gain deeper insights into the nuclei’s complex dynamics. The average migration speed of male nuclei was highly variable, with a mean front speed of 10.5 ± 16.7 µm/min (Fig 4). Male nuclei frequently exhibited back-and-forth movements. Careful analysis revealed two types of direction changes: U-turns, where the front of the nucleus executed a 180° turn and remained the front after the reversal, Reversals, where the rear of the nucleus became the new front (Fig 5B, Movies 3 & 4). In branched trichogynes, nuclei could enter one branch, change direction, and migrate into another (data not shown). Essentially, two kinds of movements were observed at both the front and rear ends of the nucleus: traction movements reminiscent of pulling forces, and retraction movements reminiscent of an elastic reaction of the nucleus. The highest traction speed recorded was 125.4 µm/min and it was for the front of the nucleus (male *H1-GFP* X female *PH-mCherry*, Fig 4, Movie S3). The highest retraction speed measured was for the rear of the nucleus: 248.9 µm/min (male *H1-mCherry* X female *β-Tubulin-GFP*, movie 5).

**Fig 5.**
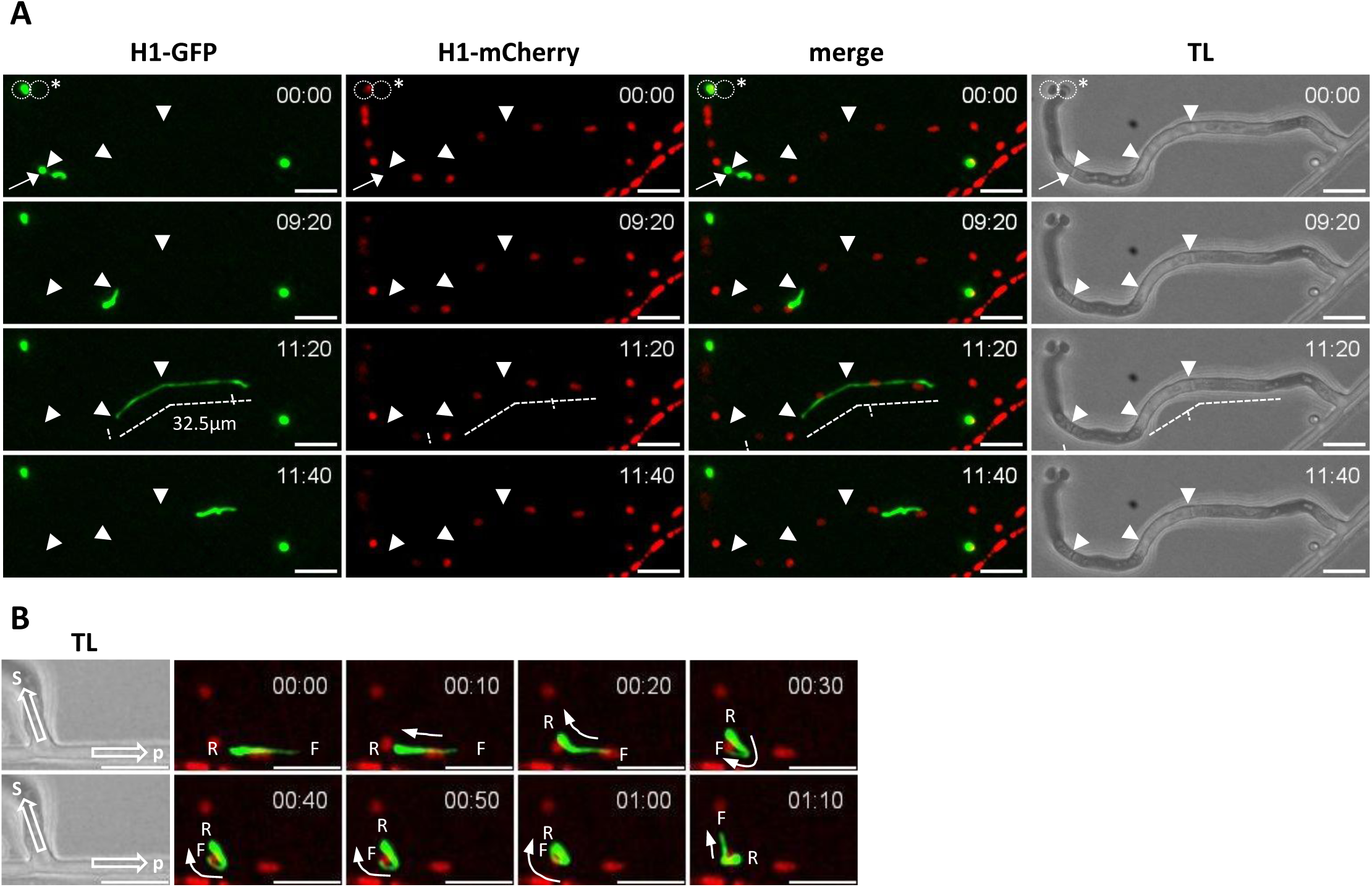
Male nuclei migration within trichogynes. Cross: male *H1-GFP* X female *H1-mCherry*. Z-projection images from time-lapse movies 3 (A) and 4 (B) acquired using a spinning disk microscope (1 frame every 10 seconds). **A)** Arrowheads indicate septa crossed by the H1-GFP-labeled male nucleus. In the T = 0 row, two spermatia are circled: the one on the left has not fused with the trichogyne and still contains its H1-GFP-labeled nucleus, while the one on the right marked with an asterisk has fused and released its nucleus, which is migrating up the trichogyne (arrow). At t = 0; the male nucleus is squeezed in the septal pore and appears split in two (but it is not). During migration, the H1-GFP-labeled nucleus exhibits back-and-forth movements and stretches while passing through the second and third septa during a single elongation phase; the maximum stretching observed is 32.5 µm (dashed line). H1-mCherry-labeled female nuclei remain stationary with only limited oscillatory movement. **B)** A migrating H1-GFP-labeled male nucleus inside the trichogyne is shown. F: front of the nucleus; R: rear of the nucleus; arrow labeled “s” indicates the direction from which the nucleus originates (spermatium), and arrow labeled “p” indicates the direction toward the protoperithecium. Arrows show the direction of nuclear movement. From t = 10 sec to t = 20 sec, the nucleus reverses direction (simple backward movement); from t = 30 sec to t = 1 min 10 sec, it performs a U-turn, with the nucleus front changing direction. TL: transmitted light. Time format: min:sec. Scale bar: 10 µm.

To further characterize the overall movement of male nuclei, we compared the speeds of the front, center, and rear in three male nuclei during stretching phases (Figs 4 & S3). Our first observation was that the mean speeds of these three regions did not differ statistically (Fig 4). Secondly, at every time point, the center speed consistently fell between the front and rear speeds (Fig S3). Therefore, when the front was being pulled forward and the rear was retracting simultaneously, the speed measured at the center could be higher than at the front. Indeed, we recorded a maximum center speed of 135.2 µm/min, surpassing the previously noted maximum front speed of 125.4 µm/min. We also compared the center speed of male nuclei during stretching and non-stretching phases, excluding pausing intervals to evaluate whether stretching correlated with faster movement. When not stretching, nuclei moved at an average speed of 10.7 ± 14.8 µm/min, whereas during stretching, their speed increased significantly to 18.3 ± 26.2 µm/min (t-test, *P* = 0.008), indicating that nuclei move faster while stretched.

Male nuclei were observed contorting and elongating when passing by female nuclei and crossing septal pores, with an average stretched length of 27 ± 17 µm (Table S1, Figs 5 & S2). The most extreme stretching measured reached 71 µm (male *H1-mCherry* X female *β-Tubulin-GFP*, Fig S2, Movie S2), approximately twelve times their relaxed length and 35 times larger than regular vegetative or female nuclei. Among the different cell organelles, resident female nuclei and large vacuoles, along with septal pores, likely represent the principal physical obstacles to migrating male nuclei. Female nuclei, with a diameter of 2.1 ± 0.2 µm, occupy most of the trichogyne cross-sectional area, which averages 3.5 ± 0.6 µm (Table S1). Trichogynes are composed of compartments separated by septa (Fig 1D). While the diameter of septal pores in *P. anserina* vegetative hyphae is about 200 nm (S. Brun, unpublished data), the diameter of septal pores in trichogynes remains unknown. Assuming a similar size, the relaxed male nuclei width of 0.8 ± 0.2 µm is roughly four times larger than the pores. Consistently, in nearly all recorded movies, male nuclei often paused before septa, slackening to relaxed form, then stretching again to pass through (Movies 3, 4, 5, 7, 8, 12 & S2). However, we also observed male nuclei crossing septal pores without pausing, sometimes crossing two septa consecutively while stretched (*e.g*., Fig 5, Movie 3 at t=11:20). This indicated that relaxation before pore passage was frequent but not obligatory. For instance, in Fig 5, the male nucleus crossed the second septum without stopping before (t = 11:20, movie 3). Together, these data demonstrate that male nuclei migration is a complex process. Male nuclei migrate rapidly and undergo extreme deformation, likely necessary to bypass bulky obstacles such as female nuclei and, most importantly, to traverse septal pores.

### Actin depolymerization does not affect nuclei movements

The major cytoskeletal components responsible for nuclear movements in eukaryotes are actin filaments and MTs. However, neither of these cytoskeletal elements has been studied in *P. anserina* to date. To investigate the actin network, we first constructed a LifeAct-GFP reporter that specifically binds filamentous actin (see Materials & Methods, Table 1) (Freitag et al., 2004; Riedl et al., 2008; Lichius and Read, 2010). We analyzed the actin network in LifeAct-GFP expressing vegetative hyphae and in *H1-mCherry LifeAct-GFP* spermatia. To study actin dynamics during fertilization, we crossed *H1-mCherry* males with *LifeAct-GFP* females and observed actin staining patterns in trichogynes before plasmogamy and during male nuclei migration after plasmogamy. Actin was present in spermatia (Fig 6A) as well as in trichogynes both before and after plasmogamy during male nuclei migration (Figs 6B, 6C; Movies 6, 7 & 8). Prior to plasmogamy, the actin network in trichogynes (Fig 6B, Movie 6) resembled that observed in vegetative hyphae (Fig 6D, Movie S4), characterized by a high concentration of actin at the trichogyne tip, visible filaments, and numerous actin patches. Filament formation was highly dynamic in both trichogynes and vegetative hyphae, consistent with previous observations in other fungi (Berepiki et al., 2010, 2011). Remarkably, the actin network became much more prominent in trichogynes during male nuclei migration. It consisted mainly of patches but was dominated by longitudinal actin cables spanning the trichogyne compartments, with a particularly high concentration at septa (Fig 6C, Movies 7 & 8). Moreover, we occasionally observed co-localization of H1-mCherry tagged, stretched male nuclei with LifeAct-GFP labeled actin cables, though this was not consistent during every nuclear stretching event (Fig 6C). To test whether the actin network was essential for male nuclei movements, we treated samples with Latrunculin B (Lat B), a drug that depolymerizes actin filaments (Spector et al., 1983), and monitored male nuclei dynamics in trichogynes. Four hours after inoculating *LifeAct-GFP* females with *H1-mCherry* spermatia (male *H1-mCherry* X female *LifeAct-GFP*), fertilized samples were mounted in 25 µM Lat B (Fig 7, Movie 9), and both actin network integrity and male nuclei movements were observed. Strikingly, complete actin depolymerization in Lat B-treated trichogynes (Fig 7A) did not affect either male nuclei speed (Fig 7B) or nuclear stretching (Fig 7C, Movie 9). This indicates that actin filaments do not play a major role in male nuclei movements and deformation within trichogynes.

**Fig 6.**
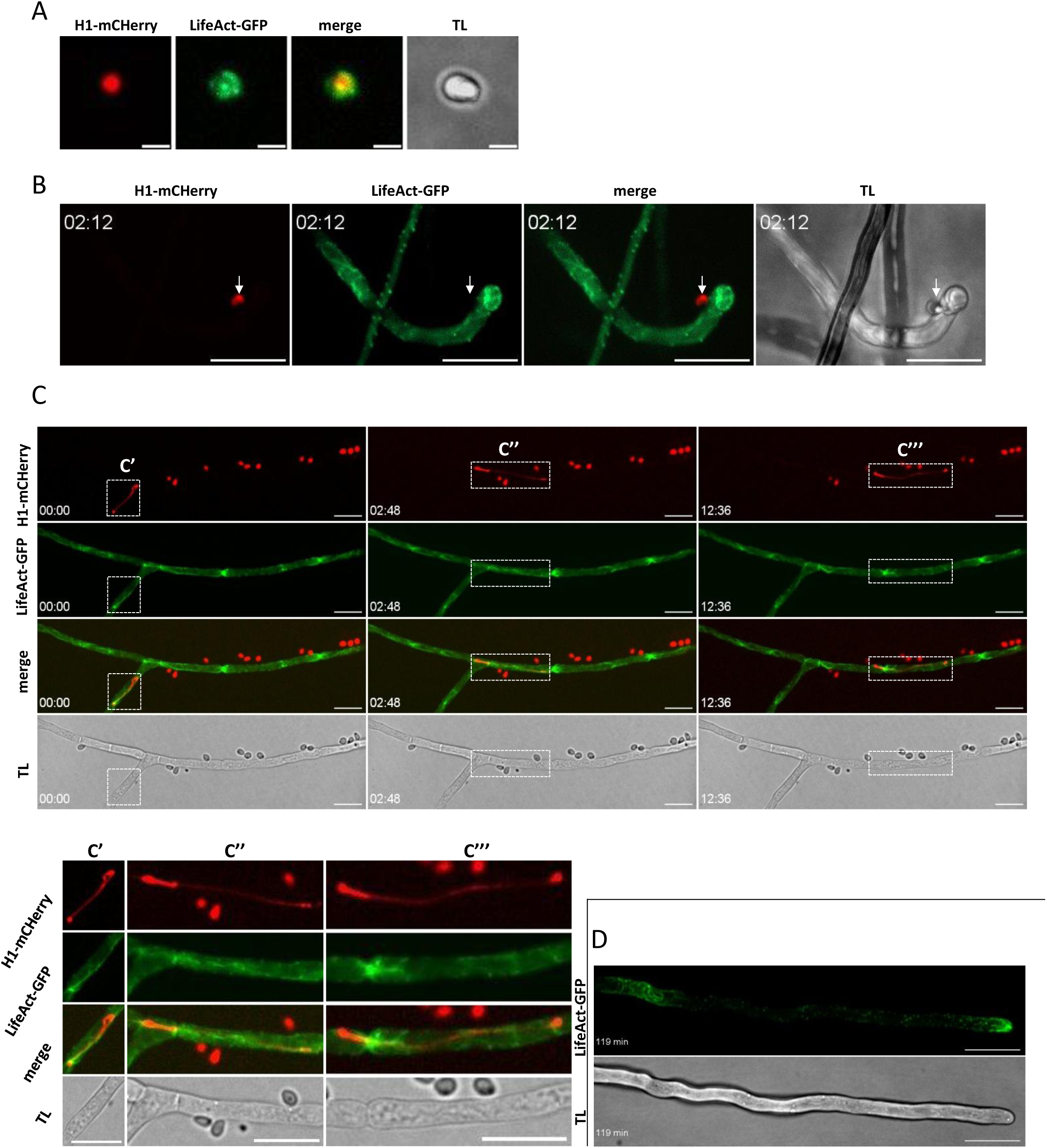
Actin cytoskeleton in spermatia, vegetative hyphae, and trichogynes. **A)** Spermatium expressing H1-mCherry and LifeAct-GFP. Z-projection images showing actin organization. Scale bar: 2 µm. **B)** Trichogyne prior to plasmogamy from a cross between male *H1-mCherry* and female *LifeAct-GFP*. Z-projection images from time-lapse movie 6 acquired using a spinning disk microscope (1 frame every 12 seconds). The H1-mCherry-labeled spermatium that attracted the trichogyne is indicated with an arrow. Time format: min:sec. Scale bar: 10 µm. **C)** Trichogyne during male nucleus migration from a cross between male *H1-mCherry* and female *LifeAct-GFP*. Z-slices from time-lapse movie 7 acquired using a spinning disk microscope (1 frame every 12 seconds). Movie 7 captures two H1-mCherry-labeled nuclei migrating sequentially within a LifeAct-GFP-labeled trichogyne. The first and second columns show the first migrating nucleus, and the third column shows the second. Panels C’, C’’, and C’’’ are high magnifications of the regions indicated by dashed rectangles. During migration, male nuclei show alternating phases of colocalization between H1-mCherry signal and LifeAct-GFP-labeled actin cables (C’, C’’), and phases where colocalization is partial or absent (C’’’). Multiple spermatia containing H1-mCherry-labeled nuclei are visible in the field of view. TL: transmitted light. Scale bar: 10 µm. D) Vegetative hyphae expressing LifeAct-GFP. Z-projection images from time-lapse movie S4 acquired using a spinning disk microscope (1 frame per minute). Scale bar: 10 µm.

**Fig 7.**
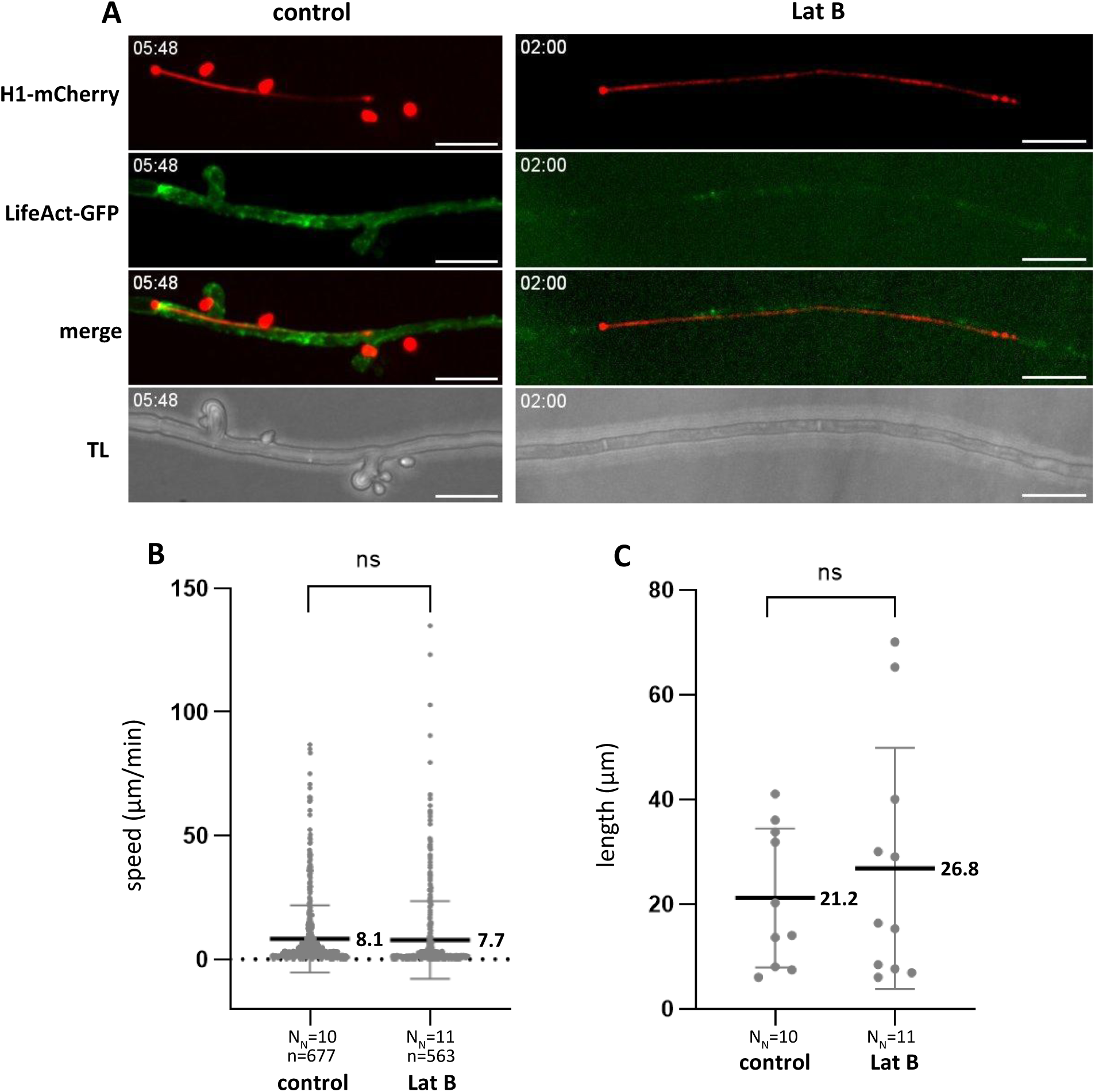
Actin depolymerization does not impair male nuclei movements and stretching. Cross: male *H1-mCherry* X female *LifeAct-GFP*, treated with 25 µM Latrunculin B (Lat B) or untreated (control). **A)** Z-projection images from time-lapse movies acquired on a spinning disk microscope. Control: movie 8 (1 frame every 12 seconds); Lat B: movie 9 (1 frame every 15 seconds). Despite Lat B-mediated depolymerization of actin filaments, male nuclei movement and stretching remain unaffected. In the control condition, four spermatia with H1-mCherry-labeled nuclei are visible within the field of view. Time format: min:sec. Scale bar: 10 µm. **B)** Male nuclei front speed. The front speed of male nuclei was measured at each time point throughout the time-lapse recordings. Mean speeds are indicated in the graph. No significant difference in speed was observed following actin depolymerization. N_N_: number of nuclei analyzed; n: number of individual front speed measurements. **C)** Maximum male nuclei stretching. For each nucleus, the greatest stretching length observed during the movie was recorded. Mean maximum stretching lengths are shown in the graph. N_N_: number of nuclei analyzed. Control was treated with 0.3% DMSO. Welch’s t-test: ns = not significant.

### MTs are polarized in trichogynes

MTs have never been studied in *P. anserina*. To investigate them, we constructed reporter strains in which the β-subunit of MT protofilaments is tagged with either GFP or mCherry (see Materials & Methods, Table 1). We analyzed the MT network in *β-Tubulin-GFP* vegetative hyphae and in *H1-mCherry β-Tubulin-GFP* spermatia. To study MT dynamics during fertilization, we crossed *H1-mCherry* males with *β-Tubulin-GFP* females and observed MT labeling in *β-Tubulin-GFP* trichogynes. The β-Tubulin-GFP labeled MT network in trichogynes was markedly different from that in vegetative hyphae. In vegetative hyphae, long longitudinal MT bundles were observed at the apex (Fig 8D’, Movie S5), consistent with observations in other species (Riquelme et al., 2018). More proximally, the MT network appeared more scattered, displaying both longitudinal bundles and bundles originating from intensely GFP-labeled, motile foci (Fig 8D’’, Movie S6). Notably, septa in vegetative hyphae showed little to no β-Tubulin-GFP signal (see below).

**Fig 8.**
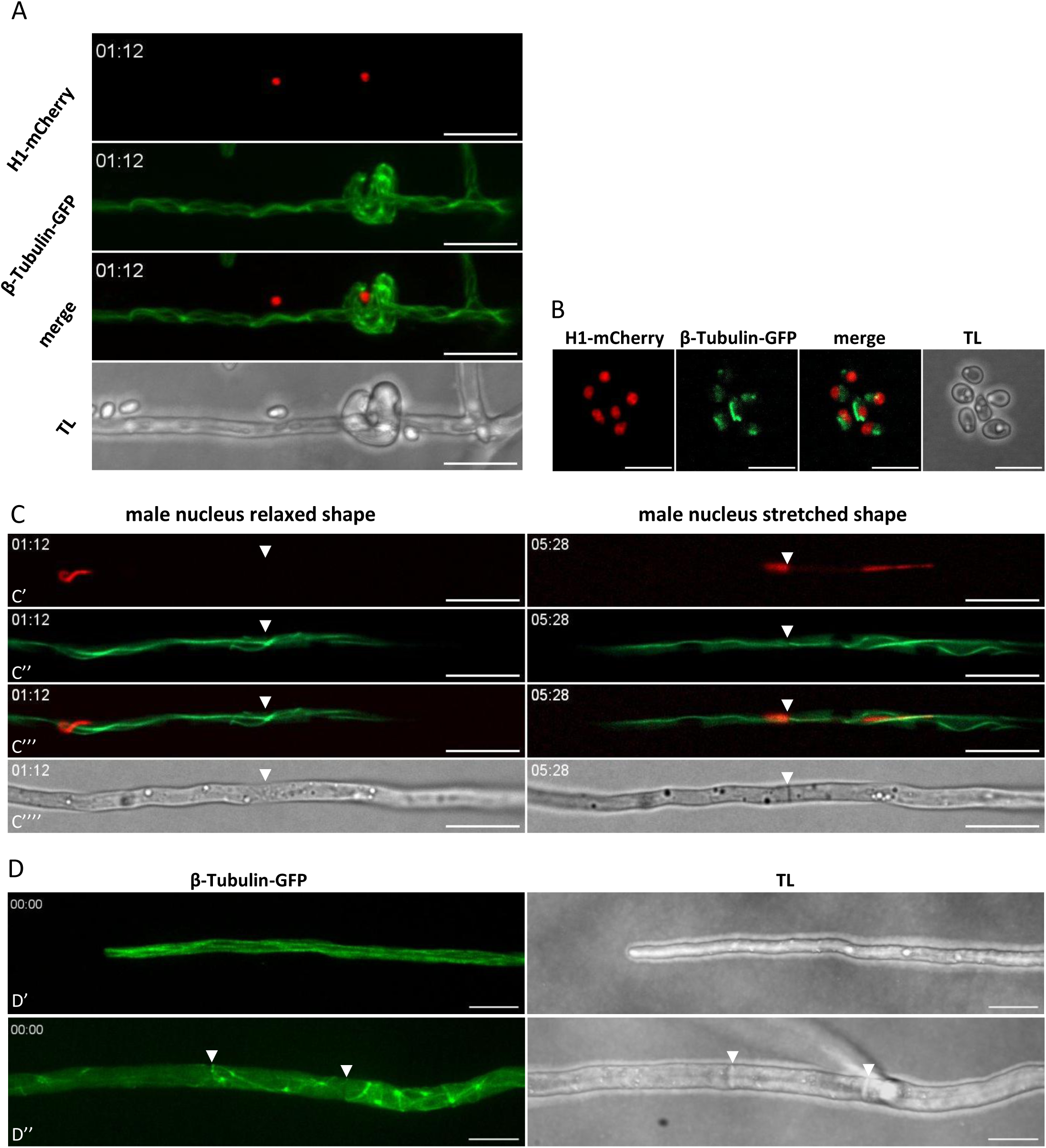
Microtubule network organization in *P. anserina*. **A)** Microtubule (MT) organization in a trichogyne prior to plasmogamy, from a cross between male *H1-mCherry* X female *β-Tubulin-GFP*. Z-projection images from time-lapse movie 10 (1 frame every 8 seconds). The spermatium on the right attracts the trichogyne, which coils around it. Time format: min:sec. Scale bar: 10 µm. **B)** MT labeling in spermatia expressing H1-mCherry and β-Tubulin-GFP. Z-projection images. Scale bar: 5 µm. **C)** MT organization in a trichogyne during male nucleus migration, from a cross between male *H1-mCherry* X female *β-Tubulin-GFP* (mounted in 1% DMSO). Z-slice images from time-lapse movie 5 (1 frame every 8 seconds). Time format: min:sec. Panels: C’, H1-mCherry channel; C’’, β-Tubulin-GFP channel; C’’’, merged channels; C’’’’, transmitted light. The migrating H1-mCherry-labeled male nucleus moves from left to right. In the left panel, the nucleus is in a relaxed shape with no visible interaction between its front and MTs. In the right panel, the front of the nucleus is being pulled, the nucleus is stretched, and its front colocalizes with MTs. Arrowhead: septum. Scale bar: 10 µm. **D)** MT organization in vegetative hyphae expressing β-Tubulin-GFP. Z-projection images from time-lapse recordings: D’, hyphal apex from movie S5; D’’, hyphal segment from movie S6. Arrowheads indicate septa. Frame rate: 1 frame every 10 seconds. Time format: min:sec. Scale bar: 10 µm.

In contrast, MTs in trichogynes predominantly consisted of thick, long bundles appearing to traverse the length of the trichogyne (Figs 8A, 8C, 9A; Movies 5, 10, 11 & 12). In fungi, spindle pole bodies (SPBs) as well as septa function as important microtubule-organizing centers (MTOCs) (Zekert et al., 2010; Zhang et al., 2017; Riquelme et al., 2018). As noted above, β-Tubulin-GFP labeling at vegetative hyphal septa was weak (Fig 8D’’). Accordingly, we rarely observed MT bundles growing from septa in vegetative hyphae: among twenty-three septa analyzed over 10-minute recordings, only one MT bundle elongation was detected (data not shown). In sharp contrast, *β-Tubulin-GFP* trichogynes exhibited intense labeling at septa along with numerous MT bundle extensions primarily oriented toward the apex (Fig 9A, Movie 11). Although tools for specifically tagging MT minus (-) or plus (+)-ends are not available in *P. anserina*, these MT bundle extensions were clearly visible, allowing us to quantify them and determine their growth direction and speed (0.17 ± 0.07 µm/s). We counted the number of MT bundle extensions at septa (within a single focal plane over 10 minutes) to calculate the percentage growing toward the apex versus toward the protoperithecium, thereby assessing MT polarity in trichogynes (Figs 9B & 9C). Because MTs can cross septal pores, the extensions observed at septa may include both MTs polymerizing at septa and those crossing the pores. First, we observed that the average number of MT bundle extensions per septum was significantly higher during male nuclei migration (10.1 ± 2.6) compared to before plasmogamy (3.8 ± 2.5). More importantly, 92 ± 16% of these MT bundles grew toward the trichogyne apex before plasmogamy, and 96 ± 6% during male nuclei migration. Taken together, these data demonstrate that the MT network in trichogynes is highly dynamic and polarized both before and during male nuclei migration. Indeed, the vast majority of MT (+)-ends extend toward the apex, which is opposite to the direction of male nuclei migration.

**Fig 9.**
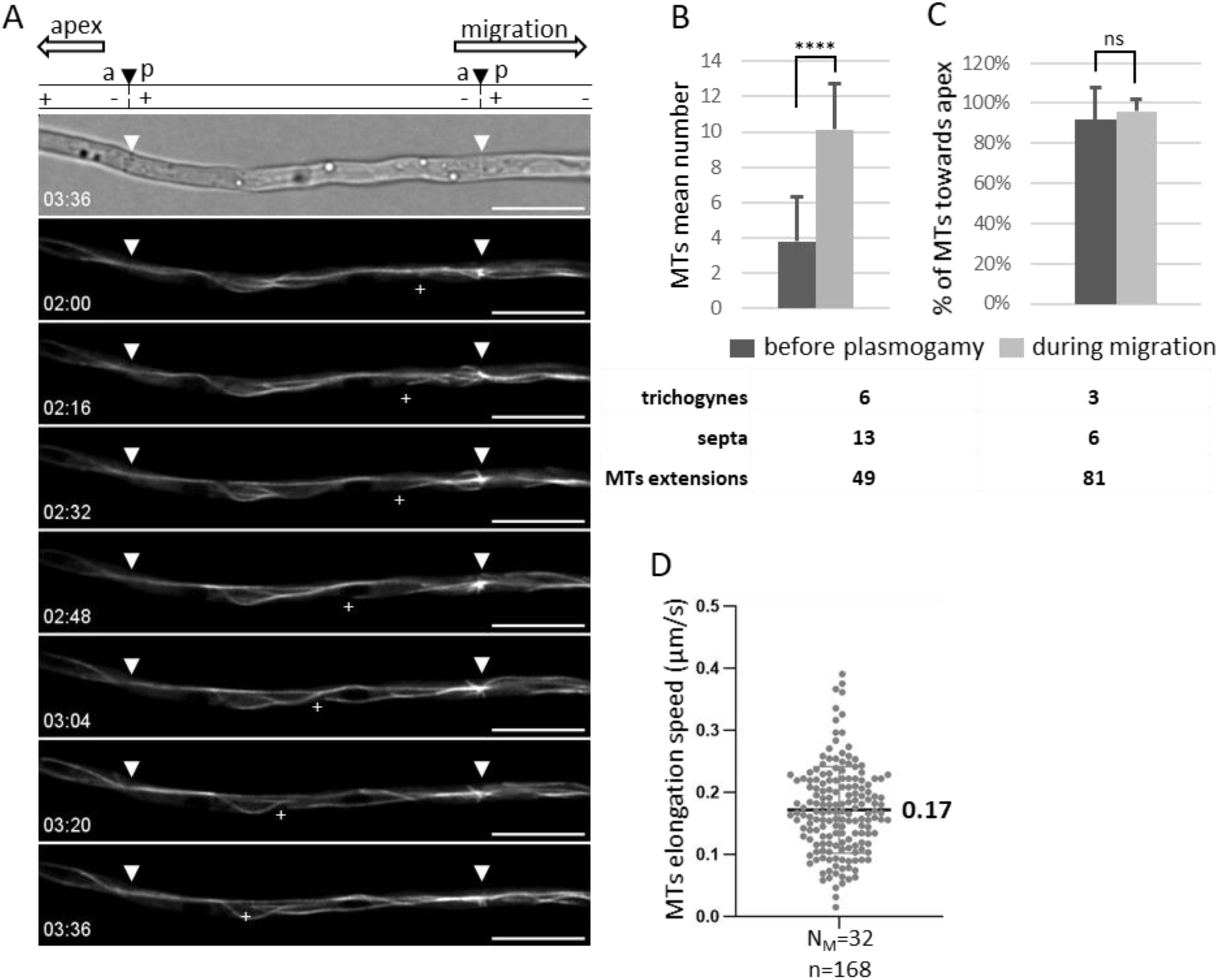
The microtubule network in trichogynes is polarized. Cross: male *H1-mCherry* X female *β-Tubulin-GFP* (mounted in 1% DMSO). **A)** Z-slice images from time-lapse movie 11 (1 frame every 8 seconds). Arrowheads indicate septa. At the top of panel A, the orientation of the trichogyne and the direction of male nuclei migration are shown. The schematic indicates septal orientation: “a” for apical side, “p” for protoperithecium side; “+” and “–” represent the main MT polarity within the trichogyne compartments as inferred from panels B and C. The first row shows transmitted light images; subsequent rows show β-Tubulin-GFP labeling. Several MT bundle elongations are visible across the time points, with the (+) end of the most prominent one indicated by a “+”. Time format: min:sec. Scale bar: 10 µm. **B)** Quantification of the mean number of elongating MTs on both sides of septa before plasmogamy and during male nuclei migration (after plasmogamy). Welch’s t-test: ns = not significant; **** P < 0.0001. **C)** Mean percentage of elongating MTs on the apical side of septa and directed toward the apex versus those directed toward the center of the thallus. No significant difference was found between pre- and post-fertilization percentages (χ²(1) = 2.23, P = 0.135). The number of trichogynes, septa, and elongating MTs analyzed before and after plasmogamy is indicated below the graphs. Counts were performed on a single z-slice over a 10-minute recording window. The majority of MTs in trichogynes grow toward the apex, establishing a polarized network as illustrated in the schematic. D) Speed of MT bundle elongation (µm/s); N_M_: number of MT bundle elongation events analyzed; n: number of individual speed measurements.

### Male nuclei movements depend of MTs

As shown above, actin depolymerization did not affect male nuclei movements. To investigate whether MTs are involved in these movements, we crossed *H1-mCherry* males with *β-Tubulin-GFP* females and analyzed the behavior of migrating male nuclei in relation to the MT network within trichogynes. We found that when H1-mCherry–tagged male nuclei were in a relaxed state (*i.e*., between stretching phases), there was no consistent or clear co-localization with GFP-tagged MTs (Fig 8C; Movies 5 & 12). In sharp contrast, during stretching phases, we observed a clear co-localization between the front part and the stretching regions of male nuclei and MT bundles (Fig 8C; Movies 5 & 12), suggesting that the leading edge of male nuclei was pulled along MT tracks. To functionally test the role of MTs in male nuclei migration, we depolymerized MTs using 350 µM Nocodazole (Noc), following the same protocol used for Lat B treatment: samples were mounted in Noc 4 hours after inoculating *β-Tubulin-GFP* female partners with *H1-mCherry* spermatia, and we then monitored both male nuclei movement and the MT network. Noc treatment led to a rapid and nearly complete depolymerization of GFP-labeled MT bundles (Fig 10; Movies 13–15). Strikingly, while 13 out of 13 nuclei migrated in untreated control *β-Tubulin-GFP* trichogynes, none of the 11 observed nuclei migrated in Noc-treated trichogynes. Moreover, these nuclei exhibited no stretching phases and maintained a constant length of 5.9 ± 1.3 µm, similar to the size of relaxed state of male nuclei (Table S1; Fig 10B). Notably, in the absence of MTs, male nuclei were not spherical but retained an irregular, loose shape. They were often found partially blocked or compressed within septal pores (Fig 10A-a; Movie 13), or arrested at the entry point of the trichogyne (Fig 10A-b; Movie 14). When free in the cytoplasm (Fig 10A-c; Movie 15), male nuclei exhibited only residual, uncoordinated movements, in stark contrast to the active migration observed in untreated samples (Fig 10A-control; Movies 5, 10 & S3). Together, these results demonstrate that MT depolymerization by Nocodazole severely impairs male nuclei movement and stretching, indicating that both migration and deformation of male nuclei within trichogynes are MT-dependent processes.

**Fig 10.**
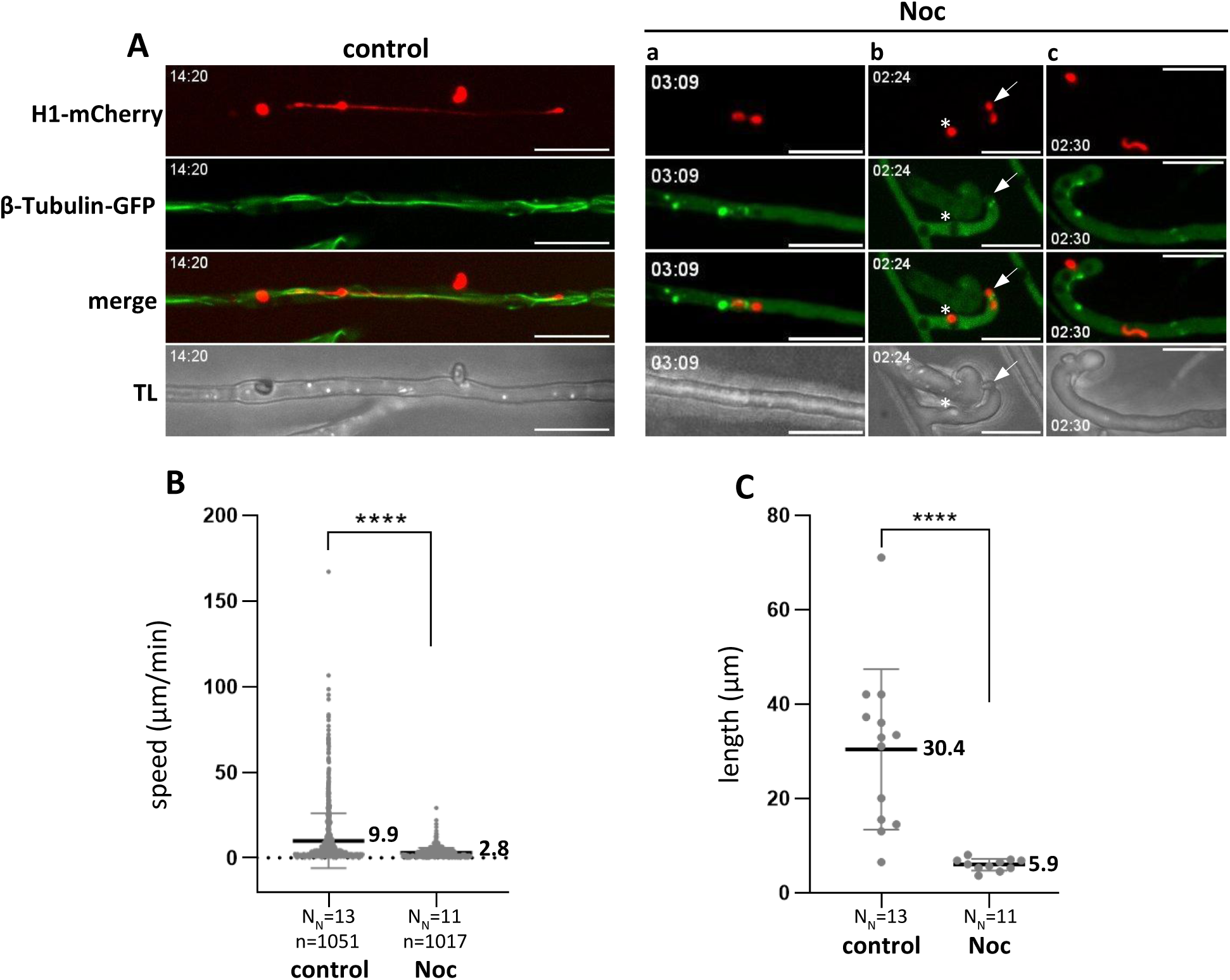
Male nuclei migration in trichogynes depends on microtubules. **A)** Z-projection images from time-lapse movies acquired on a spinning disk microscope. Comparison between untreated controls (male *H1-mCherry* X female *β-Tubulin-GFP*, mounted in 1% DMSO; movie 12, 1 frame every 10 seconds) and samples treated with 350 µM Nocodazole (Noc): **(a)** nucleus blocked at a septum (movie 13, 1 frame every 27 seconds); **(b)** nucleus blocked between the spermatium and the trichogyne (movie 14, 1 frame every 24 seconds); **(c)** nucleus floating freely in the cytoplasm (movie 15, 1 frame every 30 seconds). In the control panel, both spermatia with their H1-mCherry-labeled nuclei are visible. In panel Noc-b, an asterisk indicates an H1-mCherry-labeled nucleus remaining inside the spermatium. In panel Noc-c, a spermatium is observed contacting the trichogyne tip with its H1-mCherry-labeled nucleus still inside. MT depolymerization by Nocodazole prevents male nucleus migration and abolishes nuclear stretching. TL: transmitted light. Time: min:sec. Scale bar: 10 µm. **B)** Front speed of male nuclei. Speed was measured at each time point for all nuclei tracked in the time-lapse recordings. Mean speeds are indicated in the graph. Nocodazole treatment causes a dramatic collapse in male nuclei speed. N_N_: number of nuclei analyzed; n: number of front speed measurements. **C)** Maximum stretching of male nuclei. For each nucleus analyzed, the longest stretch observed during the recording was measured. Mean stretching lengths are shown in the graph. No nuclear stretching is observed under Nocodazole treatment. N_N_: number of nuclei analyzed. Control was treated with 1% DMSO. Welch’s t-test P-values: ns = not significant; * P < 0.05; ** P < 0.01; *** P < 0.001; **** P < 0.0001.

## Discussion

Nuclear dynamics and deformation have been studied in various model systems, including yeast, *Drosophila*, and human cells, using micropatterning approaches (Gibeaux and Knop, 2013; Golloshi et al., 2022; Lepesant et al., 2024). In this study, we explored the behavior of both female and male nuclei during sexual reproduction in the model filamentous fungus *P. anserina*, with a particular focus on male nuclei migration. The most striking observation was the remarkable ability of male nuclei to move, stretch, and contort as they migrate through the female trichogyne, mimicking inchworm movement. The conservation of this phenomenon in both *P. anserina* and *N. crassa* highlights the potential of filamentous fungi as robust models for studying extreme nuclear dynamics and deformation *in vivo*. Male nuclei migration is characterized by: i) exceptionally high migration speed, ii) dramatic stretching, particularly when navigating through obstacles such as female nuclei and septal pores, iii) the ability to perform back-and-forth movements, and iv) the capacity to select the correct path toward the protoperithecium. Furthermore, we observed that female nuclei dynamics within trichogynes are significantly reduced during male nuclei migration. This mirrors findings in *N. crassa* and suggests a conserved inhibitory effect of male nuclei migration on female nuclei motility (Brun et al., 2021).

We found that multiple male nuclei can migrate simultaneously within trichogynes, frequently leading to polyfertilization of protoperithecia. A detailed analysis of protoperithecial fertilization revealed that 50% of fertilized protoperithecia received more than one male nucleus. Upon entry into the protoperithecium, male nuclei ultimately accumulate in its central region—presumably the ascogonium—where they are expected to engage in the next stages of sexual development, including proliferation and meiosis, ultimately leading to ascospore production (Peraza-Reyes and Malagnac, 2016). In scenarios involving genetically distinct male nuclei, one would anticipate the formation of genetically mosaic perithecia producing asci of different paternal origins. However, when females were fertilized with spermatia of different genotypes, we did not observe any mosaic perithecia. We propose two hypotheses to explain this apparent discrepancy between cytological observations and genetic outcomes. First, polyfertilized protoperithecia may degenerate during development and fail to mature into perithecia capable of producing ascospores. Second, the ascogonium has not yet been specifically identified due to the lack of molecular markers, and thus we cannot definitively determine whether all male nuclei entering the protoperithecium also enter the ascogonium. It remains possible that only one male nucleus successfully enters the ascogonium, while supernumerary nuclei are excluded before the onset of the subsequent developmental stages. In support of this, it was recently shown that once fertilization has occurred, perithecia in *P. anserina* become refractory to further fertilization within a few hours (Bidard et al., 2025). Further studies will be necessary to elucidate the fate of male nuclei within the protoperithecium. Answering these questions will require significant advances in live imaging of nuclear dynamics within the protoperithecial core. Beyond nuclear proliferation, several key processes take place at this stage, including internuclear recognition prior to karyogamy and meiosis (Zickler et al., 1995), as well as genome defense mechanisms such as Repeat-Induced Point mutation (RIP) which remains poorly characterized in *P. anserina* (Gladyshev, 2017). High-resolution, live-cell imaging of these events will be crucial to uncover the spatial and temporal dynamics governing nuclear behavior and genome regulation during sexual development in filamentous fungi.

In this study, we investigated whether the actin or MT cytoskeletal networks are responsible for enabling the striking movements and deformations of male nuclei during their migration through trichogynes. Although both cytoskeletal components are more prominent in trichogynes than in vegetative hyphae, our data strongly support a predominant role for MTs in this process. While actin cables were occasionally co-localized with stretched male nuclei, the depolymerization of actin filaments using Latrunculin B did not impair male nuclei migration or stretching, arguing that the actin cytoskeleton does not play a major role in these dynamics. In contrast, MTs form extensive and dynamic bundles throughout the trichogyne, particularly at septa, and their depolymerization using Nocodazole resulted in a complete arrest of male nuclei migration and a loss of stretching behavior. This clearly demonstrates that both movement and deformation of male nuclei in trichogynes are MT-dependent. Previous studies have shown that MTs can cross septal pores (Pieuchot et al., 2015; Mouriño-Pérez et al., 2006), and while we did not directly confirm that the MT bundles observed at septa traverse these pores, we did observe male nuclei crossing multiple septal pores in a continuous, stretched state. This continuity of movement, despite the presence of septa, strongly suggests that male nuclei are pulled along MT tracks that may extend across septal pores, providing a structural and functional basis for their directed migration.

One of the most intriguing abilities of male nuclei is their capacity to orient themselves within branched trichogynes, effectively choosing the “correct path” toward the protoperithecium and reversing direction when initially taking a wrong turn at branch points. Our study provides important insights into how male nuclei may achieve this orientation by analyzing MT dynamics. In fungal cells, the main microtubule organizing centers (MTOCs) are the spindle pole bodies (SPBs) associated with nuclei and septa (Zekert et al., 2010; Zhang et al., 2017; Riquelme et al., 2018). We focused on septa due to their pronounced β-Tubulin-GFP tagging observed in trichogynes. Unlike in vegetative hyphae, where MT growth at septa is infrequent, we detected numerous MT bundle extensions at septa in trichogynes. Although specific tagging of MT (+) or (-)-ends is unavailable in *P. anserina*, the directionality of these MT bundle extensions allowed us to infer the overall MT polarity within trichogyne compartments. Our data reveal that the vast majority (>92%) of MT bundles grow towards the trichogyne apex, indicating a strong polarization with most MT (+)-ends directed apically and (-)-ends oriented toward the protoperithecium. This polarized MT network is consistent both before plasmogamy and during male nuclei migration. These observations strongly suggest that male nuclei migration is predominantly a (-)-end-directed movement along MTs toward the protoperithecium. We therefore propose that male nuclei navigate branched trichogynes by following the predominant MT (-)-end polarity, which effectively guides them retrogradely to the protoperithecium. Occasionally, the minority of MTs (∼4%) with (-)-ends oriented toward branch apices could misdirect nuclei, causing them to initially migrate in the wrong direction. In general, the ability of male nuclei to change direction at branch points or within compartments might depend on two key properties: (i) performing retrograde ((-)-end-directed) movements and (ii) switching between MT bundles oriented in opposite directions. Ultimately, the global polarization of MTs, with (-)-ends pointing toward the protoperithecium, ensures that nuclei migration is biased in the correct direction. Regarding the molecular machinery driving this movement, although microtubule-associated molecular motors (MMs) have not been studied in *P. anserina*, genomic analysis has identified putative kinesins and components of the Dynein/Dynactin complex (DDC) (Espagne et al., 2008). While kinesins capable of retrograde transport cannot be excluded, the DDC—known universally to mediate retrograde transport along MTs in eukaryotes (Riquelme et al., 2018; Reck-Peterson et al., 2018; Xiang, 2018; Singh et al., 2024)—represents the most plausible candidate driving male nuclei migration toward the protoperithecium. Conversely, kinesins responsible for anterograde transport toward MT (+)-ends may occasionally associate with nuclei, potentially explaining the observed changes in direction.

We also demonstrate that male nuclei change direction within trichogynes through two distinct mechanisms: by performing U-turns or by reversing their movement (*i.e*., backward migration). Assuming that male nuclei movements result from pulling forces generated by molecular motors (MMs) anchored to the nuclear envelope (NE) at the leading edge of the nucleus, these two modes of directional change raise important questions about the spatial distribution of such pulling forces. In the case of U-turns, the front of the nucleus changes direction, implying that a single pulling site localized at the nucleus’s front drives this reorientation. Conversely, reversing movement suggests that an additional pulling site located on the opposite, rear side of the nucleus becomes engaged to mediate backward migration. In metazoans, the LINC complex mechanically couples the NE to cytoskeletal motors, thereby transmitting forces necessary for nuclear positioning and movement. The LINC complex has been studied in both yeasts *Saccharomyces cerevisiae* and *Schizosaccharomyces pombe* where it is involved in SPB functioning (*i.e*., duplication and insertion into the NE), NE homeostasis, chromatin tethering to the NE and chromosomes movements during meiosis (Koszul et al., 2008; Wanat et al., 2008; Fan et al., 2020; Chen et al., 2019; Friederichs et al., 2011; Jaspersen, 2021; Sosa Ponce et al., 2020; Jaspersen et al., 2006; Hagan and Yanagida, 1995; Matsuda et al., 2017; Hou et al., 2012; Fernández-Álvarez et al., 2016). Whether the LINC mediates pulling forces driving male nuclei movements in trichogynes remains an important open question.

Another remarkable feature of migrating male nuclei in fungi is their exceptionally high migration speed. Because male nuclei in trichogynes are not spherical, we defined distinct regions — front, center, and rear — to accurately describe their dynamics and quantify velocity. Since nuclear movements are presumably driven by MT-associated MMs exerting pulling forces on the NE, the speed of the nuclear front, which corresponds to active traction, serves as the most relevant parameter to estimate the motor velocity. The highest front speed measured in our study was 2.1 µm/sec (125 µm/min), a value comparable to dynein-driven retrograde transport speeds reported in *Ustilago maydis* (Schuster et al., 2011), yet strikingly fast for nuclear migration in general (Cadot et al., 2015).

Another remarkable characteristic of male nuclei during migration is their exceptional flexibility. In their relaxed state, male nuclei exhibit a loose, irregular shape that is markedly different from the more compact morphology of female nuclei within trichogynes or vegetative nuclei. Upon stretching, their length can increase dramatically—from approximately 6 µm to up to 71 µm—representing nearly a twelve-fold elongation. Moreover, when traversing septal pores, male nuclei must deform significantly: they squeeze through pores estimated to be around 200 nm in diameter, despite their width being roughly 800 nm in their relaxed form, which is about four times larger than the pore size. Given the retrograde, (-)-end directed migration of male nuclei along MTs, we propose that this pronounced stretching would primarily be initiated by the binding of the DDC to the male NE, which would exert pulling forces on the nuclei front toward the MT (-)-ends. Male nuclei subjected to these MT-associated motor-driven traction forces undergo dramatic elongation, particularly as they concurrently squeeze through septal pores or pass by bulky organelles such as female nuclei. Supporting this model, we observed clear co-localization of the male nuclei front with MT bundles during the stretching phases. Conversely, we hypothesize that disruption of the proposed nucleus-DDC-MT linkage leads to relaxation of the nuclei and cessation of migration. Consistent with this, during periods when male nuclei exhibit relaxed morphology, no co-localization with MT bundles at male nuclei extremities was detected.

The migration of male nuclei is a long-range process during which they repeatedly squeeze through septal pores and undergo significant stretching, thereby facing intense and repetitive mechanical stress. Remarkably, we have never observed any male nucleus rupturing during migration, indicating that this process is highly robust and suggesting that male nuclei are structurally adapted to withstand such mechanical challenges. One of the clearest adaptation is their loose, irregular shape, which contrasts sharply with the round or ovoid morphology typical of vegetative and female nuclei. Importantly, this loose nuclear shape appears independent of the actin and MT cytoskeletons, as male nuclei retain their characteristic morphology even when both cytoskeletal networks are depolymerized. This suggests that this unique nuclear flexibility is an inherent feature of the structure and composition of the male nucleus itself. In metazoans, lamins play a central role in determining nuclear stiffness and shape; however, lamins are absent in fungi, and to our knowledge, no other nuclear components have been demonstrated to fulfill a similar structural role in fungal nuclei (Davidson and Lammerding, 2014; Meseroll and Cohen-Fix, 2016). While the lack of lamins might partially explain the looseness of fungal nuclei in general, it does not clarify why this loose shape is specific to male nuclei and not observed in vegetative or female nuclei. We propose that various factors may contribute to the enhanced flexibility of male nuclei, including differences in NE composition— both lipids and proteins—as well as chromatin organization and its potential tethering to the NE (Janota et al., 2020; Burla et al., 2020). Future work aimed at identifying the molecular determinants of male nuclei flexibility and robustness will provide valuable insights into nuclear mechanotransduction and adaptation to mechanical stress.

Movements and deformations of the fungal nucleus during sexual reproduction are remarkably complex, representing some of the most extreme nuclear dynamics described across all eukaryotes. A lot of work remains to be done before to fully characterize these processes. We propose that studying mechanotransduction in male nuclei during sexual reproduction in the fungus *Podospora anserina* offers a powerful model to investigate the molecular mechanisms enabling their exceptionally rapid movement. This research could reveal how nuclei endure extreme mechanical stresses and uncover the potential impacts of these forces on chromatin organization and gene expression.

## Materials & Methods

### Strains (Table 1) and culture conditions

The strains used in this study are all listed in Table 1. All *P. anserina* strains derive from the wild-type S strain, ensuring a homogeneous genetic background (Espagne et al., 2008; Rizet, 1952). The *PaPks1^193^* mutant, deficient for the polyketide synthase-encoding gene acting at the first step of melanin synthesis, is described in (Coppin and Silar, 2007). Standard culture conditions, media, and genetic methods for *P. anserina* were described in (Rizet and Engelmann, 1949; Silar, 2013) and on the *Podospora anserina* database http://podospora.i2bc.paris-saclay.fr/. The M0 medium is similar to M2 medium but without dextrin.

### Strains construction

#### H1-GFP & H1-mCherry

The genomic sequence of the putative H1 histone (*Pa_5_8740*), including its 854 bp upstream regulatory region, was amplified using the Phusion polymerase (https://www.thermofisher.com) with primers H1-TAGF (5’-CTCGAGCAGCTTAGCGGCAGATCGAG-3’), bearing an *X*hoI site at the 5’ end, and H1-TAGR (5’-AAGCTTCGCCGAGGCAGCTTCCGCCT-3’), bearing a *H*indIII site at the 5’ end, and omitting the stop codon of *Pa_5_8740* CDS. 3’-OH thymidines were added in a subsequent step with the GoTaq polymerase to enable cloning into pGEM-T (https://france.promega.com/). The PCR product was cloned into pGEM-T, digested with *X*hoI and *H*indIII (FastDigest, https://www.thermofisher.com), and inserted into pBC-hygR-eGFP or pBC-hygR-mCherry digested with the same enzymes, creating the pBC-hygR-H1-GFP and pBC-hygR-H1-mCherry transformation vectors conferring hygromycin B resistance [hygR]. Both vectors were transformed into the wild-type *S* strain, and two independent [hygR] primary transformants per construct were selected based on nuclear fluorescence intensity. Subsequently, the respective *H1-GFP* and *H1-mCherry* primary transformants (named so for simplicity) were backcrossed with the *S* strain, and homokaryotic progeny of each mating type (*mat+* and *mat-*) were selected. Only one *H1-mCherry* and one *H1-GFP* transformant were used in this study.

#### β-Tubulin-GFP & β-Tubulin-mCherry

Both constructs, composed as follows — <*S*peI site – 444 bp regulatory sequence of the *AS4* gene (strong, constitutive *AS4* promoter (Silar et al., 2001; Lacaze et al., 2015)) – GCCACC (Kozak sequence) – *Pa_4_7630* CDS (putative β-Tubulin) without stop codon – *eGFP* or *mCherry* CDS – *S*peI site> — were synthesized by IDT (https://eu.idtdna.com/). Both pUCIDT-β-Tubulin-GFP and pUCIDT-β-Tubulin-mCherry were digested by *S*peI (FastDigest, https://www.thermofisher.com), and the inserts were subcloned into pBC-phleoR (conferring phleomycin resistance) digested with *S*peI, generating the pBC-phleoR-β-Tubulin-GFP and pBC-phleoR-β-Tubulin-mCherry transformation vectors. These vectors were transformed into the wild-type *S* strain, and two independent [phleoR] primary transformants per construct were selected based on MT network fluorescence intensity. The primary transformants were backcrossed with the *S* strain, and homokaryotic progeny of each mating type (*mat+* and *mat-*) were selected. These strains were named *β-Tubulin-GFP* and *β-Tubulin-mCherry* for simplicity. Only one *β-Tubulin-GFP* transformant was used in this study. *β-Tubulin-GFP* was crossed with *H1-mCherry*, and [hygR phleoR] *H1-mCherry β-Tubulin-GFP* double-tagged homokaryotic progeny of each mating type were selected.

#### LifeAct-GFP

Two long reverse complementary oligonucleotides of the following sequence: AAGCTTGCCACCATGGGCGTCGC<u>C</u>GACCTCATCAAGAAGTTCGAGTCCATCTCCA AGGAGGAG<u>G</u>GGGCCC, composed as follows — <*Hi*ndIII site – GCCACC (Kozak sequence) – *Neurospora crassa* Lifeact sequence (Berepiki et al., 2010) encoding the Lifeact 17 AA actin-binding peptide (MGVADLIKKFESISKEE) (Riedl et al., 2008) with the codon GCT>A changed to GCC>A (underlined) – additional G (underlined) for in-frame cloning – *A*paI site> — were synthesized by IDT (https://eu.idtdna.com/). A mix of both oligonucleotides was used for direct 3’-OH thymidine addition by GoTaq (https://france.promega.com/) followed by cloning into pGEM-T (https://france.promega.com/). pGEM-T-LifeAct-GFP was digested by *H*indIII and *A*paI (FastDigest, https://www.thermofisher.com), and the insert was subcloned into the pBC-hygR-AS4-GFP transformation vector digested by *H*indIII and *A*paI (Lacaze et al., 2015). In the pBC-hygR-AS4-LifeAct-GFP vector, transcription of the LifeAct-GFP reporter is driven by the strong constitutive *AS4* promoter. This vector was transformed into the wild-type *S* strain, and one [hygR] primary transformant was selected based on actin network fluorescence intensity. The primary transformant was backcrossed with the *S* strain, and homokaryotic progeny of each mating type (*mat+* and *mat-*) were selected. The strain was named *LifeAct-GFP* for simplicity. *LifeAct-GFP* was crossed with *H1-mCherry*, and [hygR] homokaryotic *H1-mCherry LifeAct-GFP* progeny of each mating type, exhibiting both nuclei (red) and actin (green) tagging, were selected.

#### PH-mCherry

The AS4-PH-mCherry construct, composed as follows — <*S*peI site – 444 bp *AS4* promoter – GCCACC (Kozak sequence) – 552 bp *Homo sapiens* Pleckstrin-homology (PH) plasma membrane phosphoinositide-binding domain (Várnai and Balla, 1998) – mCherry CDS – *S*peI site> — was synthesized and cloned into pUCIDT(Amp) by IDT (https://eu.idtdna.com/). The pUCIDT-AS4-PH-mCherry vector was digested with *S*peI (FastDigest, https://www.thermofisher.com), and the insert subcloned into the pBC-nouR transformation vector (conferring nourseothricin resistance) digested with *S*peI. pBC-nouR-AS4-PH-mCherry was transformed into the wild-type *S* strain, and two [nouR] primary transformants were selected based on plasma membrane fluorescence intensity. These transformants were backcrossed with the *S* strain, and homokaryotic progeny of each mating type (*mat+* and *mat-*) were selected. The strain was named *PH-mCherry* for simplicity. Only one *PH-mCherry* strain was used in this study. This strain was crossed with *H1-GFP*, and homokaryotic [hygR nouR] *H1-GFP PH-mCherry* progeny of each mating type were selected.

#### NLS-GFP & NLS-mCherry

Both AS4-NLS-GFP and AS4-NLS-mCherry constructs, composed as follows — <*H*indIII site – 444 bp *AS4* promoter – GCCACC ATG (Kozak sequence + start codon) – type one NLS (Nuclear Localization Sequence, 84 bp encoding PDNTPPRKRSKVSRACDECRRKKIKCDA) of the putative transcription factor *Pa_4_9410* (S. Brun, unpublished) – *eGFP* or *mCherry* CDS – *H*indIII site> — were synthesized and cloned into pUCIDT(Amp). These inserts were digested by *H*indIII (FastDigest, https://www.thermofisher.com) and subcloned into pBC-genR (conferring geneticin/G418 resistance) and pBC-phleoR, respectively, generating pBC-genR-AS4-NLS-GFP and pBC-phleoR-AS4-NLS-mCherry transformation vectors. pBC-genR-AS4-NLS-GFP was transformed into the wild-type *S* strain, and two [genR] primary transformants were selected based on nuclear fluorescence intensity. The strain was named *NLS-GFP* for simplicity. The *NLS-GFP* primary transformants were backcrossed with the *S* strain, and homokaryotic [genR] progeny of each mating type was selected. Only one *NLS-GFP* strain was used in this study. *NLS-GFP* was crossed with *H1-mCherry*, and [hygR genR] homokaryotic *H1-mCherry NLS-GFP* double-tagged progeny of each mating type were selected.Similarly, pBC-phleoR-AS4-NLS-mCherry was transformed into the wild-type *S* strain. Two [phleoR] primary transformants were selected based on nuclear fluorescence intensity. The strain was named *NLS-mCherry* for simplicity. The *NLS-mCherry* primary transformants were backcrossed with the *S* strain, and homokaryotic [phleoR] progeny of each mating type was selected. Only one *NLS-mCherry* strain was used in this study. *NLS-mCherry* was crossed with *H1-GFP*, and [hygR phleoR] homokaryotic *H1-GFP NLS-mCherry* double-tagged progeny of each mating type were selected.

### Live-Cell-Imaging

#### Vegetative hyphae live imaging

The strains analyzed were grown on solid M2 medium, and samples were mounted inverted in water in 2-well µ-Slide ibidiTreat chambers (https://ibidi.com/). Images were acquired using a motorized inverted microscope Zeiss spinning disk CSU-X1, as described below.

Fertilization procedure. Petri dishes containing solid M0 medium were inoculated with the “female” strain (*mat-* or *mat+*) and incubated for 5–7 days to allow protoperithecia development. The inoculum was placed on a 3 × 3 cm piece of 3M paper (https://www.3mfrance.fr) serving as a carbon source, and observations were performed on the mycelium growing in the translucent M0 medium. Under these conditions, mycelium density is reduced, which facilitates microscopic observation while allowing sexual development to proceed. A 2–3 cm² piece of M0 medium containing protoperithecia was cut and transferred to an empty Petri dish. This sample was inoculated with 100–200 µL of a 1–2 × 10⁶ spermatia/mL suspension of the “male” partner of the opposite mating type, in sterile H₂O with 0.01% Tween 20. Spermatia suspensions were prepared on the day of inoculation.

#### Fertilization live imaging

After 4 hours of incubation at 27 °C in a moist chamber, the sample was mounted inverted in one of the following conditions: sterile water, 350 µM Nocodazole, 1% DMSO (Nocodazole control), 25 µM Latrunculin B, or 0.3% DMSO (Latrunculin B control), using a 2-well µ-Slide ibidiTreat chamber. Samples were then inspected for fertilization events. Images were acquired using a Zeiss spinning disk CSU-X1 motorized inverted microscope (Oberkochen, Germany), equipped with four lasers (405 nm, 488 nm, 561 nm, 640 nm) for fluorescence imaging and their respective filters, and a sCMOS PRIME95 camera (Photometrics), at the ImagoSeine Imaging Facility: https://www.ijm.fr/plateformes-et-plateaux-techniques/imagoseine/. For low-magnification imaging (×25 and ×40 objectives), z-stacks covering a 50 µm range were acquired and processed into z-projection images. Time-lapse acquisition was carried out over 10–15 hours with a frame rate of 1 image every 3–5 minutes. For high-magnification imaging (×63 objective), time-lapse acquisition was performed over 5–20 minutes with a frame rate of 1 image every 6–24 seconds and a variable z-range of 8–25 µm. Z-stacks were processed into z-projection images, except for co-localization analyses. Images were analyzed with FIJI (Schindelin et al., 2012), except for quantification of female nuclei movements, which was performed using IMARIS (https://imaris.oxinst.com/).

#### Male nuclei speed measurements

The speed of the front, center, and rear of male nuclei was measured manually at each time point of the time-lapse recordings. Front speeds were most frequently analyzed in this study. The highest front traction speed determined was 125 µm/min, based on image-by-image analysis. For each time point, we verified whether the measured front speed (only values above 100 µm/min were considered significant) corresponded to a traction or retraction movement. Rear speeds were analyzed similarly, and the highest rear speed recorded was 248 µm/min, corresponding to a nuclear retraction. For Nocodazole-treated nuclei that were blocked during migration without stretching, front speed was measured by selecting one extremity of the nucleus. For male and female nuclei speed analyses, the number of nuclei analyzed (N_N_) and the number of individual speed measurements (n) are indicated in the figures. For statistical analysis, speed values measured from different nuclei under the same condition were pooled. Conditions were compared using Welch’s t-test. p-values are indicated as follows: ns, not significant; * p < 0.05; ** p < 0.01; *** p < 0.001; **** p < 0.0001.

#### MTs elongation quantification

MT polarization in vegetative hyphae and trichogynes, both before and after plasmogamy, was determined by counting visible MT elongations at septa. For quantification, we analyzed the most β-Tubulin-GFP-tagged focal plane (corresponding to the most central z-section) and recorded MT elongations live over a 10-minute period. In total, 23 septa were analyzed in vegetative hyphae, 13 in trichogynes before plasmogamy, and 6 in trichogynes after plasmogamy during male nuclei migration. For each septum, the number of MT elongations was counted, and the mean number is presented in Fig 9B. MT elongations directed toward the apex and those directed toward the protoperithecium were counted separately. The percentage of elongations oriented toward the apex in trichogynes is also presented in Fig 9B.

## Authors’s contribution

MCV: Investigation; ZK: Investigation; PG: Investigation & Writing -review & editing; CL: Formal analysis & Writing -review & editing; AG: (group leader) Writing - review and editing. SB: Project leader: Conceptualization, Investigation, Methodology, Validation, Visualization, Writing original draft, review & editing, Project administration, Funding acquisition.

## Acknowledgments

We thank the ImagoSeine facility at the Institut Jacques Monod, a member of the France BioImaging infrastructure supported by the French National Research Agency (ANR-10-INSB-04, “Investissements d’Avenir”), and in particular Nicolas Moisan, who developed the Python script used to automatically convert the z-stacks generated during our time-lapse acquisitions into z-projections. We are also grateful to Patricia Moussounda (Institut Jacques Monod) for her valuable assistance in preparing the various media used in this study. We thank Sandra Claret from the Polarity & Morphogenesis team (Institut Jacques Monod) for providing the PH domain sequence used to construct the PH-mCherry strain. We are especially thankful to Prof. Fabienne Malagnac (Fungal Epigenomics and Development, I2BC – Université Paris-Saclay) for welcoming us into her laboratory at a time when the transformation procedure for *P. anserina* had not yet been implemented in the Polarity and Morphogenesis lab (Institut Jacques Monod). This study was supported by IdEx “Emergence” Université Paris Cité (ANR-18-IDEX-0001).

## Supplemental materials

**Fig S1. Polyfertilization of protoperithecia. A)** Fertilized perithecia from a cross between male *H1-mCHerry* X female *wild-type*. A total of 23 protoperithecia were imaged for 15 hours (1 image every 5 minutes), and the number of male nuclei entering each protoperithecium was counted. Paternal contribution in mature perithecia was assessed by fertilizing wild-type *S* females with a mixture of wild-type *S* and *PaPks1^193^*mutant spermatia. A total of 158 mature perithecia were dissected, and the ascus composition was determined. “wt” refers to perithecia containing only asci from *S* X *S* crosses (as shown in B’), “*PaPks1^193^*” indicates perithecia with asci exclusively from *PaPks1^193^*× *S* crosses (as shown in B’’), and “mosaic” corresponds to perithecia containing a mix of both types of asci. **B)** Rosettes of asci from *S* X *S* crosses (B’) and *PaPks1^193^*X *S* crosses (B’’). In *P. anserina*, most asci contain four heterokaryotic ascospores. In *S* X *S* crosses, rosettes consist of asci with four melanized *PaPks1^+^/PaPks1^+^*ascospores. In *PaPks1^193^* X *S* crosses, rosettes contain asci with two melanized *PaPks1^+^/PaPks1^+^*ascospores and two non-melanized *PaPks1^193^/PaPks^193^* ascospores. Scale bar: 100 µm.

**Fig S2. Longest observed stretching of a male nucleus.** Cross between male *H1-mCherry* and female *β-Tubulin-GFP*. Z-projection images from time-lapse movie S2 recorded using a spinning disk microscope (frame rate: 1 image every 8 seconds). The maximum length observed for the stretched male nucleus is 71 µm. TL: transmitted light. Scale bar: 10 µm.

**Fig S3. Comparison of front, center, and rear speeds during male nuclei stretching.** Speeds are given in µm/min for each analyzed time point, restricted to intervals corresponding to nuclear stretching phases. A) male *H1-GFP* X female *PH-mCherry* (movie S3); B) male *H1-mCherry* X female *LifeAct-GFP* (movie 8); **C)** male *H1-mCherry* X female *β-Tubulin-GFP* (movie 5). The asterisk (*) highlights the highest center speed recorded in the study: 135.2 µm/min.

