## supplemental figures and table for "The fastest movements and the most extreme deformations of the cell nucleus are driven by microtubules in the model fungus *Podospora anserina*"

### Slide 1
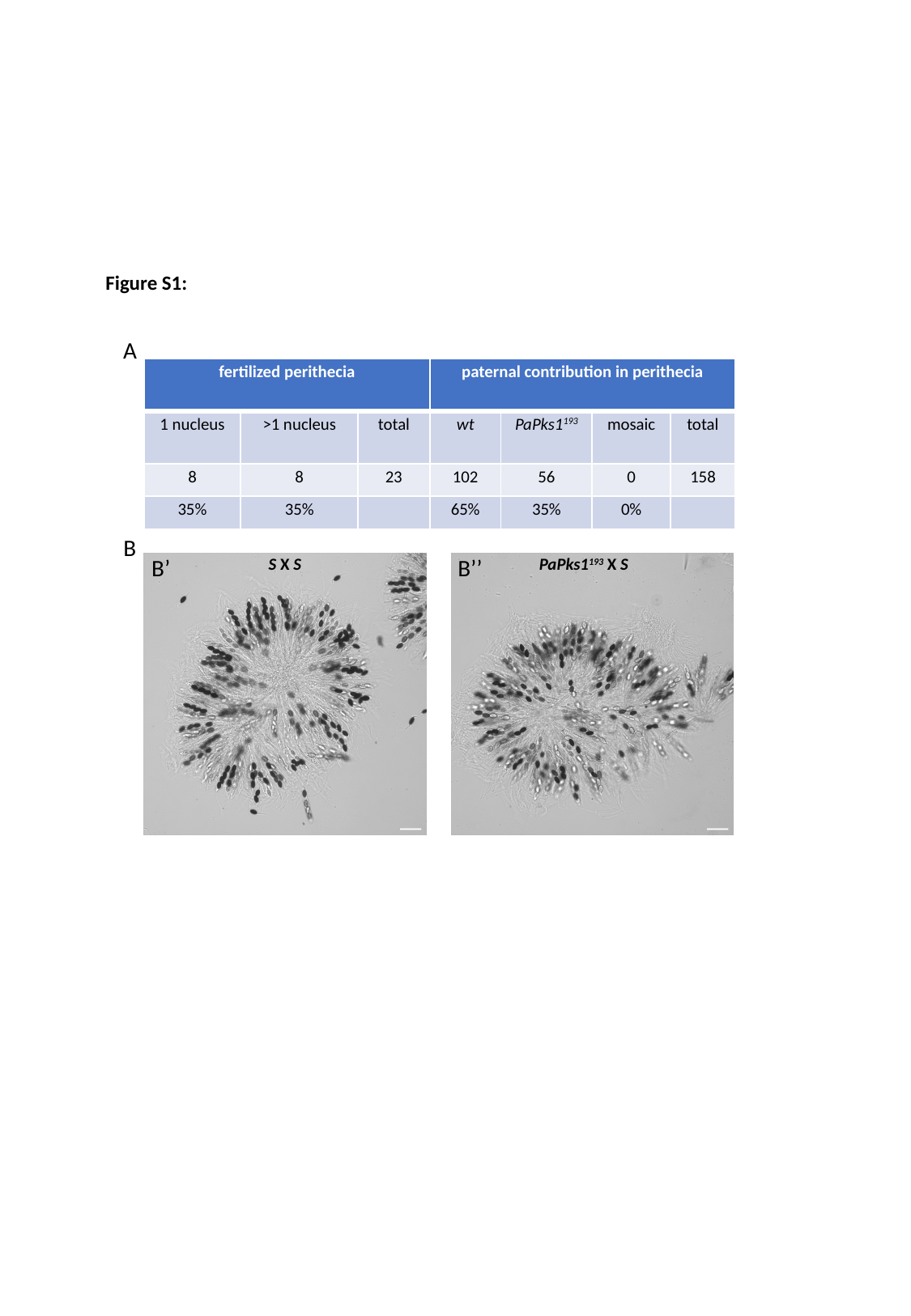

Figure S1:
A
| fertilized perithecia | | | paternal contribution in perithecia | | | |
| --- | --- | --- | --- | --- | --- | --- |
| 1 nucleus | >1 nucleus | total | wt | PaPks1^193^ | mosaic | total |
| 8 | 8 | 23 | 102 | 56 | 0 | 158 |
| 35% | 35% | | 65% | 35% | 0% | |
B
B’
B’’
S X S
PaPks1^193^ X S

### Slide 2
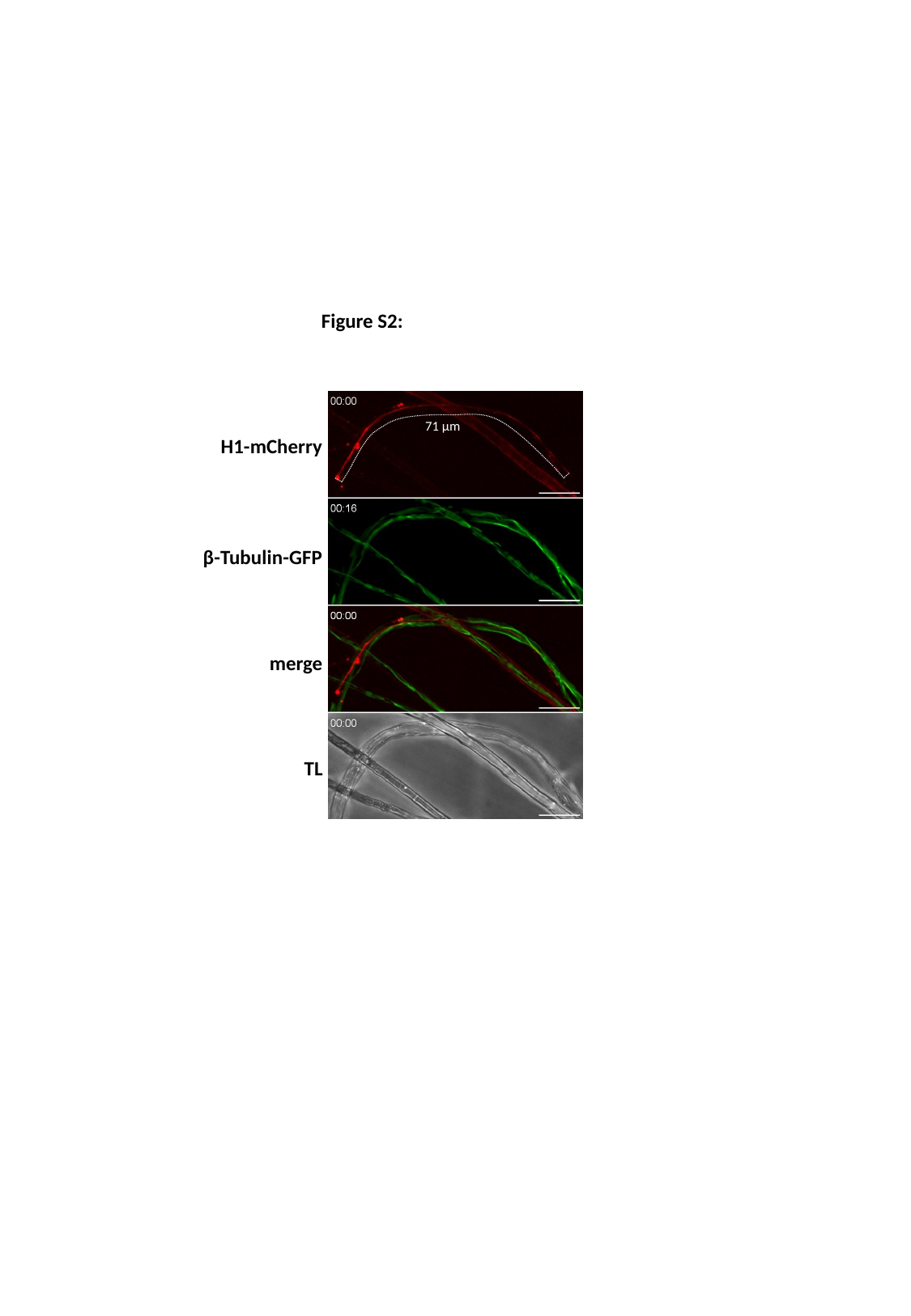

Figure S2:
71 µm
H1-mCherry
β-Tubulin-GFP
merge
TL

### Slide 3
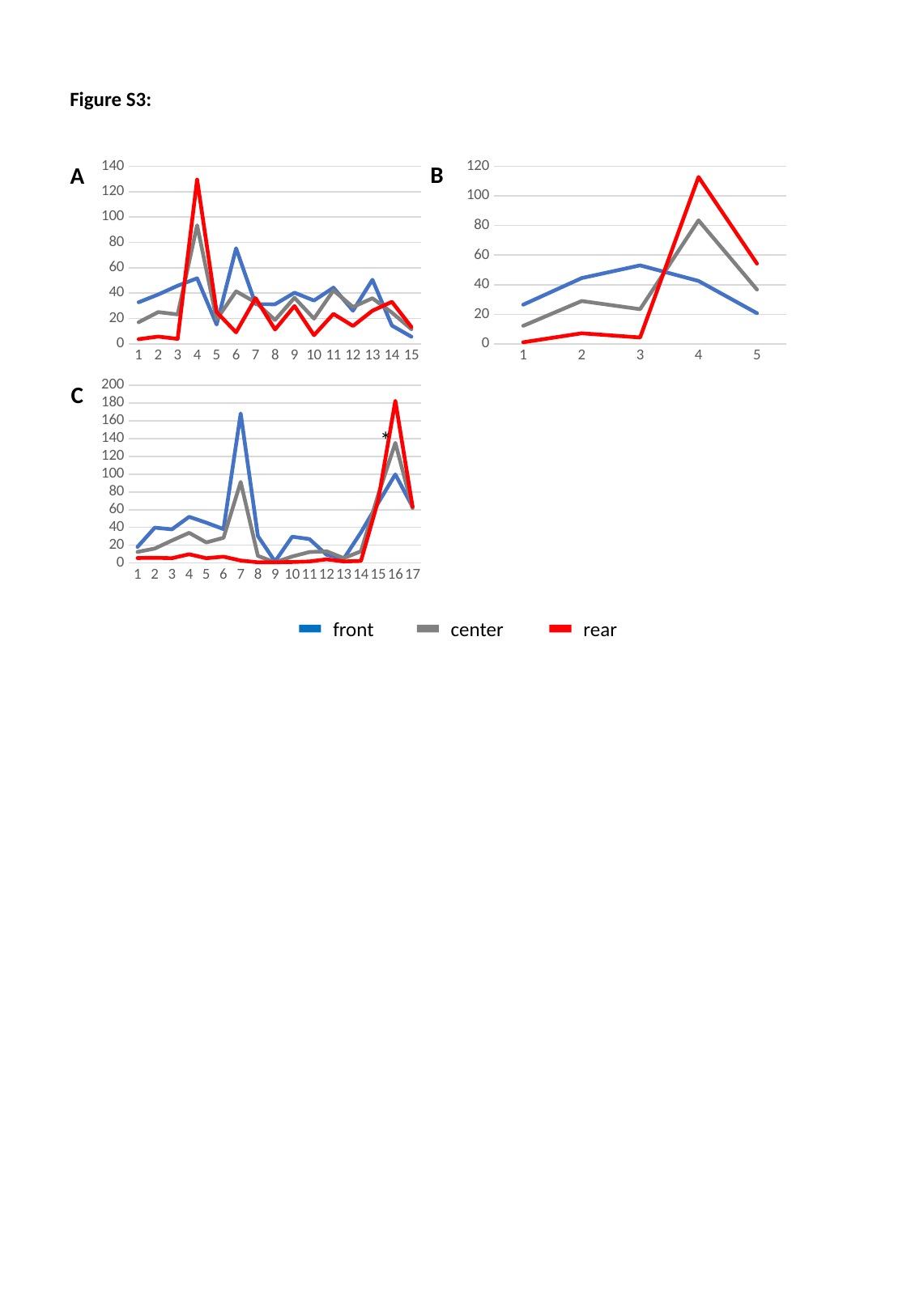

Figure S3:
B
A
#### Chart
| Category |
|---|
#### Chart
| Category |
|---|
#### Chart
| Category | | | |
|---|---|---|---|C
*
front
center
rear

### Slide 4
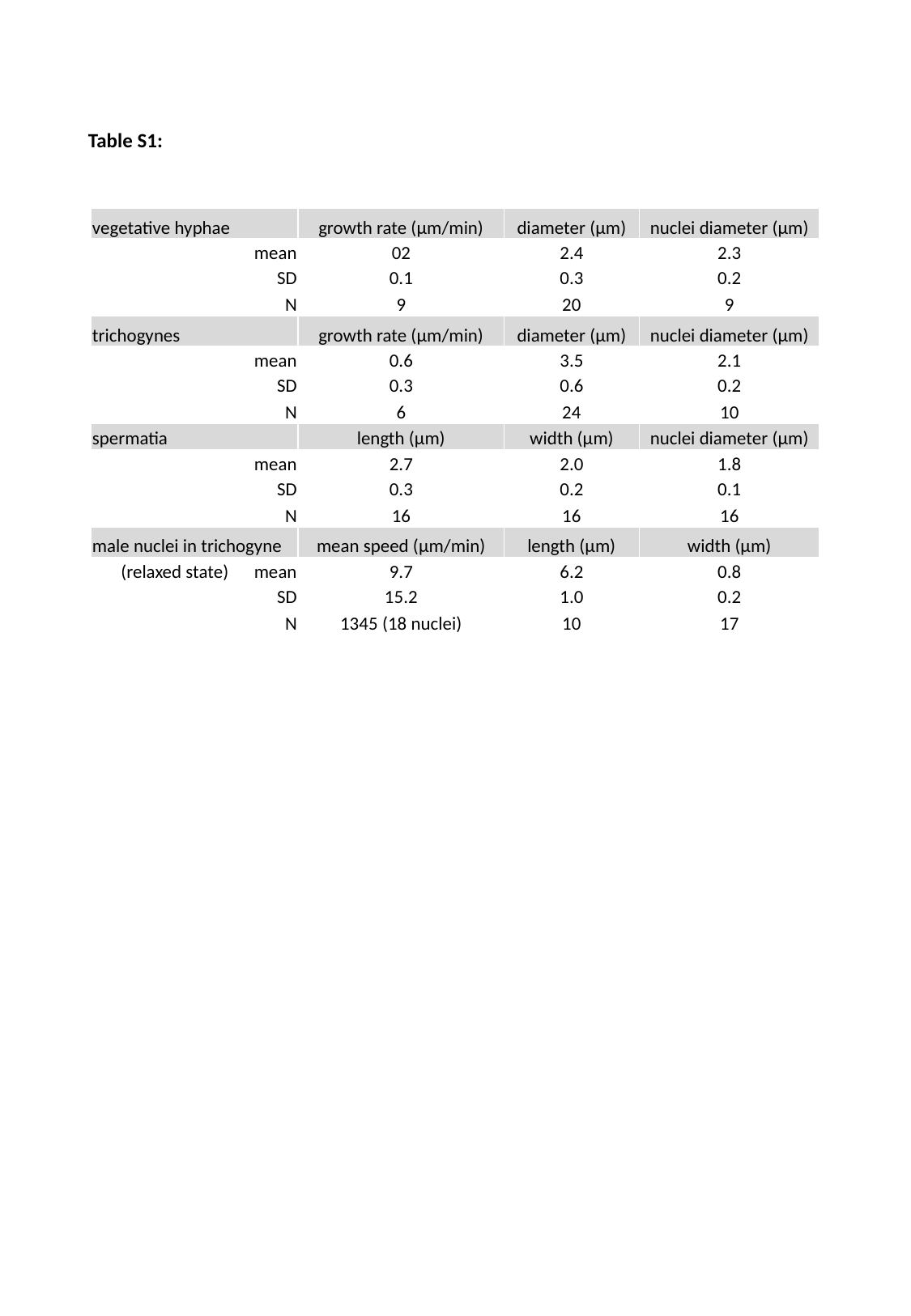

Table S1:
| vegetative hyphae | growth rate (µm/min) | diameter (µm) | nuclei diameter (µm) |
| --- | --- | --- | --- |
| mean | 02 | 2.4 | 2.3 |
| SD | 0.1 | 0.3 | 0.2 |
| N | 9 | 20 | 9 |
| trichogynes | growth rate (µm/min) | diameter (µm) | nuclei diameter (µm) |
| mean | 0.6 | 3.5 | 2.1 |
| SD | 0.3 | 0.6 | 0.2 |
| N | 6 | 24 | 10 |
| spermatia | length (µm) | width (µm) | nuclei diameter (µm) |
| mean | 2.7 | 2.0 | 1.8 |
| SD | 0.3 | 0.2 | 0.1 |
| N | 16 | 16 | 16 |
| male nuclei in trichogyne | mean speed (µm/min) | length (µm) | width (µm) |
| (relaxed state) mean | 9.7 | 6.2 | 0.8 |
| SD | 15.2 | 1.0 | 0.2 |
| N | 1345 (18 nuclei) | 10 | 17 |
